# EpiBrain: the brain’s epigenetic landscape in a snapshot

**DOI:** 10.64898/2026.01.22.701139

**Authors:** Maja Johnson, Aiden F Eno, Karl Stuntzner-Gibson, James W MacDonald, Theo K Bammler, Yijie Geng

## Abstract

Epigenetics connect nature with nurture and can help explain how genes, experiences, and the environment influence gene expression to promote individual differences in behavior and mental health. However, how these perturbations induce epigenetic changes across various regions of the brain remains largely unknown. To visualize changes in brain’s epigenetic landscape at a glance, we develop EpiBrain, a method utilizing whole-brain imaging and registration to capture epigenetic changes in the entire zebrafish brain in a snapshot. Using this method, we uncovered brain-wide epigenetic changes induced by brain activity, chemical exposures, and genetic mutations. We found that *gadd45b*, an immediate early gene with epigenetic modulatory activity, mediates experience-induced epigenetic changes in the brain, and that these changes contribute to brain activity and behavioral alterations. EpiBrain enables rapid and unbiased assessment of whole-brain epigenetic changes following physiological and disease-relevant perturbations and facilitates mechanistic discoveries at the molecular level.

## INTRODUCTION

Epigenetic mechanisms such as DNA methylation and histone modifications have been implicated in the regulations of brain function and behavior. In eusocial insects like honeybees and ants, individuals share similar genes but develop different forms and functions (social castes) due to environmentally influenced changes in gene expression, a process mediated by epigenetic mechanisms including DNA methylation and histone acetylation^1–4^. Similarly, in mammals and other vertebrate animals, epigenetic regulations are found to be pivotal for brain plasticity, learning, and behavior^5–10^. In mature neurons, epigenetic processes constantly shape synaptic plasticity and neuronal circuits, which are critical for cognition, learning, and memory^11,12^. Epigenetic dysregulations are also linked to neurodevelopmental and neuropsychiatric disorders including schizophrenia, autism, depression, and addiction, and likely underlie gene-environment interactions contributing to mental illness^13–19^.

Many environmental factors including toxicants, drugs of addiction, and experience, as well as innate factors such as genetic mutations influence an individual’s epigenetic state. For example, caste determinations in eusocial insects are controlled primarily by environmental inputs, e.g., nutrition, experience, and pheromones^2^, rather than genetics. In mammals, exposure to drugs of addiction has been found to persistently change brain’s epigenetic state, coinciding with altered behavior and neuroplasticity^20–22^. Early life experiences such as stress induce long-lasting epigenetic changes in the brain, altering behavioral outcomes and increasing susceptibility to psychopathology^23–26^. In the adult human brain, acute psychosocial stress alters DNA methylation in key neuronal genes including oxytocin receptor (*OXTR*) and brain-derived neurotrophic factor (*BDNF*)^27^. Besides environmental factors, many mental illness risk genes are epigenetic regulators. For example, mutations in the chromatin remodelers *ARID1B*, *CHD8*, and the DNA methylation enzyme *DNMT3A*, are linked to autism spectrum disorder^28,29^.

These past research suggest that epigenetics likely play a pivotal role in the regulation of brain function and behavior in health and diseases by bridging genes with environmental exposures and neuronal activities, thereby connecting nature with nurture^30–32^. Research on the neural and genetic basis of brain function in the past decades has benefited tremendously from advancements in experimental methods. Technical innovations such as optogenetics, calcium imaging, and functional magnetic resonance imaging (fMRI) have promoted unprecedented depth of investigations into the neural mechanisms underlying brain function and behavior. On the other hand, robust genetic and genomic approaches including CRISPR/Cas9 gene editing and genome-wide association studies (GWAS) have helped identify hundreds of genes associated with brain function and diseases such as autism^28,29^. In the meantime, however, how environmental factors, experiences, and genes regulate brain function and behavior through epigenetic mechanisms have remained largely unknown, mainly due to the limitations of current research methodologies and the lack of a robust experimental platform enabling systematic detection of brain-wide epigenetic changes following perturbations and stimulations.

To bridge this critical gap, we introduce EpiBrain, a method integrating deep epigenetic immunohistochemistry, whole-brain volumetric imaging, and brain registration to map the brain’s epigenetic landscape in a snapshot. EpiBrain enables systematic characterization of brain-wide epigenetic modifications triggered by diverse stimuli, including neural activity, chemical exposures, and genetic perturbations. Using EpiBrain, we identified the immediate early gene (IEG) *gadd45b* as a mediator of hindbrain DNA demethylation and reduced locomotion following chemically induced seizure-like activity. These results demonstrate EpiBrain as a powerful tool for rapid assessment of brain-wide epigenetic changes and provide mechanistic insights into how these changes mediate physiological and disease-relevant processes. Future applications of EpiBrain, including its extension to mammalian systems, will facilitate deeper understanding of the epigenetic underpinnings of brain function, behavior, and disease.

## RESULTS

### Developing the EpiBrain method

Development of the EpiBrain method was inspired by a whole-brain activity mapping method called zBrain^33^, which detects neuronal activity by comparing the relative abundance of phosphorylated extracellular regulated kinase (pErk; a neuronal activity marker elevated upon neuronal firing) with total Erk (tErk; remains constant during neuronal firing). EpiBrain adopts a similar strategy to detect changes in epigenetic marks by comparing variable epigenetic modifications with a corresponding unmodified reference signal from DNA or histone. For example, to detect changes in histone modification, larval zebrafish brains were counter-stained with two anti-histone antibodies: one recognizing an unmodified histone subunit, such as H3, and the other targeting a modification on the same subunit, such as H3K27me3. Similarly, to detect changes in DNA modifications such as 5-methylcytosine (5mC) and 5-hydroxymethylcytosine (5hmC), we counter-stained the brain with a DNA dye and an antibody detecting methylated DNA, such as anti-5mC and anti-5hmC. Following whole-brain confocal volumetric imaging, 3D brain images were registered to a reference brain to enable voxel-wise comparison of the relative signal strength of histone modifications over total histone, or DNA methylations over total DNA, between an experimental group and a control group, thereby revealing epigenetic changes induced by experimental perturbations. Together, this method provides a “snapshot” of the zebrafish brain by identifying regions where a specific epigenetic modification is upregulated or downregulated (Figure 1A).

**Figure 1.**
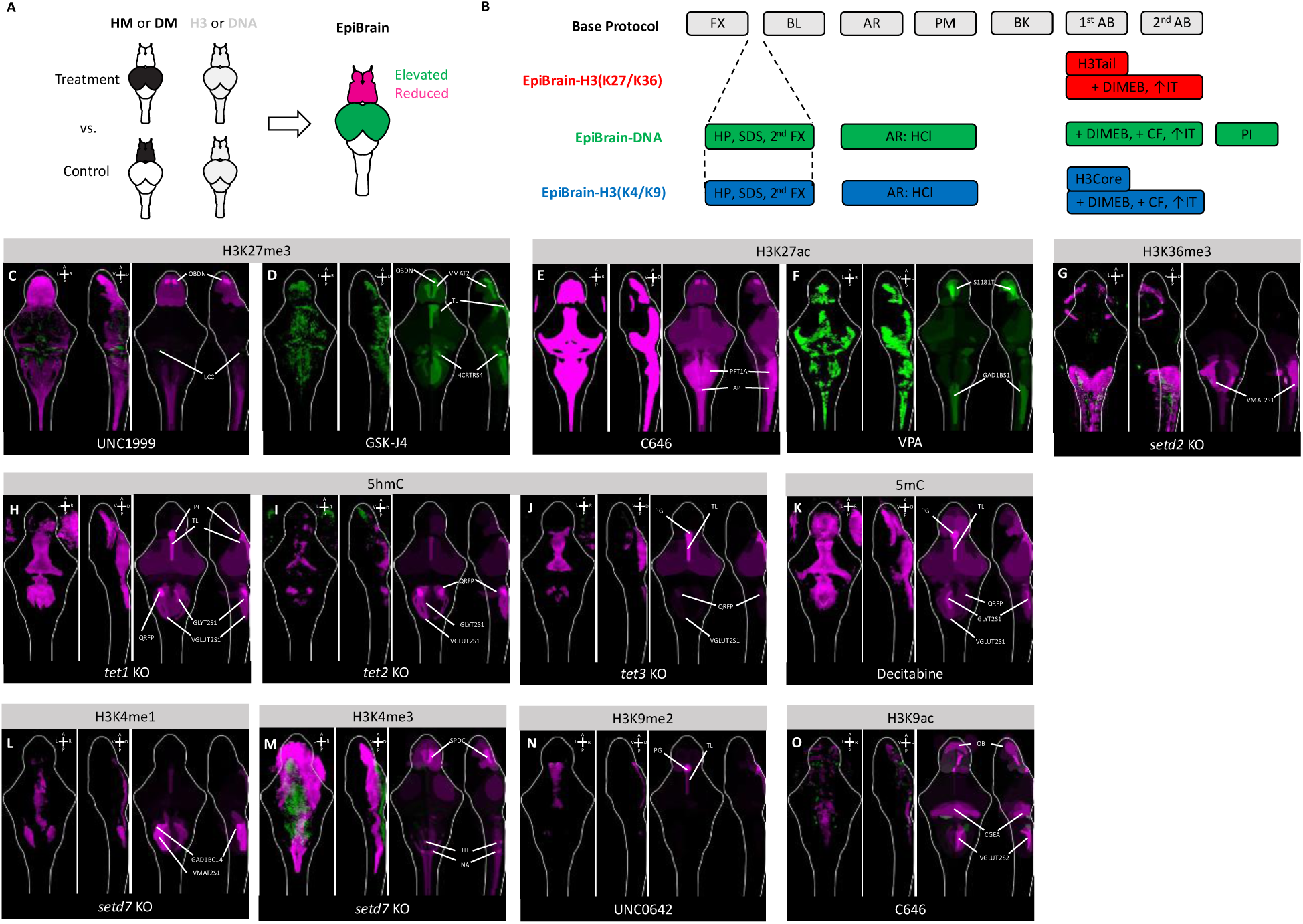
Developing the EpiBrain method. (**A**) A schematic illustration of the EpiBrain imaging and signal detection approach. Larval zebrafish are counterstained for histone modification (HM) and histone H3 (H3) or DNA modification (DM) and DNA. Whole-brain images taken from control and treatment groups are shown with differential levels of abundance for HM and DM in distinct brain regions. Images are registered to a reference brain for signal detection. Brain regions with elevated HM/DM in the treatment group as compared to the control group are labeled in green, whereas regions with reduced HM/DM in the treatment group are labeled in magenta. (**B**) Schematic illustrations of three EpiBrain staining protocols. All three protocols were developed based on a base protocol which consists of the following steps: fixation (FX), bleaching (BL), antigen retrieval (AR), permeabilization (PM), blocking (BK), primary antibody incubation (1^st^ Ab), and secondary antibody incubation (2^nd^ AB). The EpiBrain-H3(K27/K36) protocol counterstains modifications on the H3 tail that are close to the H3 core (i.e., lysine residuals K27 and K36) against H3. H3 is detected using an anti-H3-tail antibody (H3Tail) that targets the N-terminus of H3. Different from the base protocol, DIMEB is added during primary and secondary antibody incubations to boost antibody penetration. A longer incubation time (↑IT) is also applied to improve antibody binding. The EpiBrain-DNA protocol counterstains DNA modifications (i.e., 5mC and 5hmC) against the DNA. Several additional treatment steps are added between the FX and BL steps of the base protocol, including hydrogel polymerization (HP), SDS treatment, and a post-fixation step (2^nd^ FX). The AR and PM steps in the base protocol are replaced by a HCl-based antigen retrieval step (AR: HCl). Similar to EpiBrain-H3(K27/K36), DIMEB and a longer incubation time are applied to boost antibody penetration. In addition, samples are centrifuged and incubated at 37°C (CF) during primary and secondary antibody incubations to further improve antibody staining. Propidium Iodide (PI) is applied after antibody incubations to label DNA. The EpiBrain-H3(K4/K9) protocol counterstains modifications on the N-terminus of the histone H3 tail (i.e., lysine residuals K4 and K9) against H3. H3 is detected using an anti-H3-core antibody targeting the C-terminus of H3. This protocol is otherwise similar to the EpiBrain-DNA protocol minus the PI staining step. (**C-G**) EpiBrain-H3(K27/K36) staining and imaging analysis detected expected changes in the zebrafish brain’s histone modification state following targeted perturbations. The level of H3K27me3 in the brain is globally reduced by the Ezh2 inhibitor UNC1999 (C) and elevated by the H3K27me3-demethylase inhibitor GSK-J4 (D). H3K27ac level is reduced by the histone acetyltransferase p300 inhibitor C646 (E) and increased by the HDAC inhibitor VPA (F). H3K36me3 level is reduced by knocking out (KO) the histone methyltransferase *setd2* (G). Each panel consist of three images. The two images on the left are dorsal and lateral views of the brain, with changes in the relative abundance of an epigenetic modification shown in colors: green represents elevation compared to the control, and magenta represents reduction. A: anterior. P Posterior. L: left. R: right. V: ventral. D: dorsal. The image on the right highlights brain regions with significant changes in the epigenetic modification. These brain regions are labeled in colors, with green representing regions with elevated levels of epigenetic modification compared to the control and magenta representing reduced levels of modification. The brightness of the color correlates with the degree of significance. (**H-K**) EpiBrain-DNA staining and imaging analysis detects expected changes in the zebrafish brain’s DNA methylation state following targeted perturbations. The level of 5hmC is reduced by knocking out the 5mC dioxygenases *tet1* (H), *tet2* (I), and *tet3* (J). The level of 5mC is reduced by the DNMT inhibitor decitabine (K). (**L-O**) EpiBrain-H3(K4/K9) staining and imaging analysis detects expected changes in the zebrafish brain’s histone modification state following targeted perturbations. H3K4me1 level is reduced by knocking out the lysine methyltransferase *setd7* (L). H3K4me3 level is also largely reduced by *setd7* knockout, but with an elevation in the dorsal medial region (M). H3K9me2 level is reduced by the lysine methyltransferases G9a and GLP inhibitor UNC0642 (N). H3K9ac level is reduced by C646 (O). Brain regions are identified based on alignment with the zBrain atlas following brain registration. The top differentially regulated brain regions are labeled in the images. Raw analysis results for all differentially regulated brain regions and their signal intensities in images shown in Figure 1 are summarized in Supplementary Data 1. AP: area postrema. GAD1BS1: gad1b stripe 1 (spinal cord). GAD1BC14: gad1b cluster 14. GLYT2S1: glyt2 stripe 1. HCRTRS4: 6.7FDhcrtR-Gal4 stripe 4. LCC: locus caudalis cerebelli. NA: noradrendergic neurons of the interfascicular and vagal areas. OBDN: olfactory bulb dopaminergic neuron. PG: pineal gland. PFT1A: pft1a stripe. QRFP: qrfp neuron cluster sparse. S1181t: s1181t cluster. SPDC: subpallial dopaminergic cluster. TH: small cluster of TH-stained neurons. TL: torus longitudinalis. VGLUT2S1: vglut2 stripe 1 (rhombencephalon). VMAT2: vmat2 cluster (telencephalon). VMAT2S1: vmat2 stripe1 (rhombencephalon).

To develop EpiBrain, we first attempted to establish an optimal staining protocol for histone H3. Initial attempts followed a base protocol adapted from the zBrain method^33^ with modifications. Briefly, larvae at 6 days post fertilization (6 dpf) were fixed overnight and bleached to remove pigmentation. An antigen retrieval step and a permeabilization step were then performed to optimize antibody penetration, followed by blocking, primary antibody incubation, and secondary antibody incubation prior to imaging (Figure 1B).

Likely due to the presence of the high abundance of histone H3, initial attempts using a standard antibody dilution did not yield robust staining, as evidenced by poor signal intensity across the brain (Supplementary Figure 1A). Increasing the antibody concentration from 1:500 to 3:500 significantly improved the H3 signal, but still left the central brain regions under-stained (Supplementary Figure 1B). To further improve antibody penetration, we added heptakis(2,6-di-O-methyl)-β-cyclodextrin (DIMEB), a potent enhancer of cholesterol extraction and membrane permeabilization previously shown to promote deep penetration of standard antibodies into mouse tissues during whole-body immunolabeling^34^. We also extended the primary and secondary antibody incubation times to 3 days. The addition of DIMEB during the permeabilization and antibody incubation steps markedly enhanced antibody penetration and enabled homogeneous staining throughout the larval zebrafish brain (Supplementary Figure 1C).

To enable brain registration, a wild-type brain was counterstained for histone H3 and tErk (Supplementary Figure 2A) and registered to the zBrain atlas’s tErk template brain. The H3 channel of the registered image was thereafter used as the histone H3 reference template for registering EpiBrain images (Supplementary Figure 2B). We successfully counter-stained histone H3 with the histone H3 modifications lysine 27 trimethylation (H3K27me3; Supplementary Figure 3A), lysine 27 acetylation (H3K27ac; Supplementary Figure 3B), and lysine 36 trimethylation (H3K36me3; Supplementary Figure 3C), using the same optimized protocol established for histone H3 staining. This protocol is hereafter referred to as EpiBrain-H3(K27/K36) (Figure 1B).

To verify that the EpiBrain-H3(K27/K36) method can accurately detect histone H3 modifications, we applied chemical and genetic perturbations with well-characterized epigenetic modulatory activities on histone H3 lysine 27 and 36 to zebrafish larvae prior to staining. We first applied the compound UNC1999, an inhibitor of the polycomb repressive complex 2 (PRC2)’s core catalytic subunit, enhancer of zeste homolog 2 (Ezh2). As PRC2 is responsible for the methylation of histone 3 lysine 27, the PRC2 inhibitor UNC1999 is expected to cause a global reduction of H3K27me3. Indeed, we observed a pervasive reduction of H3K27me3 throughout the zebrafish brain following embryonic treatment with 10 µM UNC1999 from 0 to 3 days post fertilization (0-3 dpf) (Figure 1C). A particularly strong reduction was detected in the olfactory bulb dopaminergic neurons (OBDN) in the forebrain. Sporadic elevations of H3K27me3 were detected in the medial midbrain and rostral hindbrain regions, such as in the hindbrain locus caudalis cerebelli (LCC), likely reflecting compensatory mechanisms occurring after UNC1999 withdrawal (3 dpf) and prior to EpiBrain staining (6 dpf). In addition, exposure to GSK-J4^35^, a dual inhibitor of the H3K27 lysine demethylases KDM6A (UTX) and KDM6B (JMJD3), at 40 µM from 0 to 6 dpf resulted in robust brain-wide upregulation of H3K27me3 (Figure 1D), including in the forebrain OBDN and vesicular monoamine transporter 2 (vmat2) neuronal clusters (VMAT2), the midbrain torus longitudinalis (TL), and the hindbrain hypocretin receptor neurons 6.7FDhcrtR-Gal4 stripe 4 (HCRTRS4) and adjacent hindbrain regions.

Similarly, for H3K27ac, a pronounced downregulation was observed across the brain following exposure to the histone acetyltransferase p300 inhibitor C646 (Figure 1E), with particularly strong downregulations in the hindbrain pancreas transcription factor 1α (ptf1a) neurons (PTF1A) and area postrema (AP), whereas a pervasive brain-wide upregulation of H3K27ac was detected following embryonic exposure to the histone deacetylase (HDAC) inhibitor valproic acid (VPA) (Figure 1F), with notable upregulations in the forebrain s1181t enhancer trapped neurons (S1181T) and hindbrain glutamate decarboxylase 1b (gad1b) neuronal stripe 1 (GAD1BS1).

The SET domain-containing protein 2 (*SETD2*) gene encodes a histone methyltransferase that catalyzes trimethylation of histone H3 at lysine 36 (H3K36me3). Interestingly, knocking out its zebrafish ortholog, *setd2*, selectively suppressed H3K36me3 in the hindbrain vmat2 stripe 1 (VMAT2S1) as well as in muscles surrounding the eyes and spinal cord (Figure 1G).

The EpiBrain-H3(K27/K36) staining procedure was unsuccessful when applied to detect DNA methylations 5-methylcytosine (5mC) and 5-hydroxymethylcytosine (5hmC) (Supplementary Figure 4A). Based on prior reports, we reasoned that more stringent denaturing conditions were required to adequately expose DNA antigens for antibody binding. We therefore tested several denaturing approaches, including heat treatment at 80 °C, heating in sodium citrate buffer^36,37^, and hydrogen chloride (HCl), and found that HCl treatment most effectively enhanced the staining signal (Supplementary Figure 4B).

However, HCl treatment also caused substantial distortion of brain morphology (Supplementary Figure 4B), rendering accurate registration to the reference brain difficult. To better preserve tissue architecture during denaturation, we embedded larval zebrafish in an acrylamide tissue hydrogel prior to HCl treatment using an adaptation of the passive CLARITY technique (PACT)^38^. A sodium dodecyl sulfate (SDS)-based delipidation step was incorporated to achieve sample clearing and permeabilization during the PACT procedure. Hydrogel embedding markedly improved tissue morphology following HCl treatment, yet antibody penetration remained suboptimal (Supplementary Figure 4C). Finally, by applying centrifugal pressure, a method previously shown to enhance antibody penetration in thick tissues^39^, we ultimately achieved deep and homogeneous penetration of both anti-5mC and anti-5hmC antibodies, as evidenced by the improved staining signal (Supplementary Figure 4D).

We next tested several commonly used DNA dyes, including DAPI and propidium iodide (PI), for compatibility with this staining procedure. While DAPI was not compatible with HCl denaturation in our protocol (data not shown), consistent with a previous report^40^, PI (Supplementary Figure 5A) showed excellent compatibility with 5mC/5hmC co-staining (Supplementary Figure 5B-5C) and was therefore used as the reference DNA stain for subsequent experiments. Similar to the H3 reference brain, we created a PI template brain by registering a tErk/PI counterstained brain (Supplementary Figure 5D) to the tErk template brain (Supplementary Figure 5E). We hereafter refer to this protocol as EpiBrain-DNA (Figure 1B).

To verify the effectiveness of the EpiBrain-DNA protocol, we first knocked out the zebrafish Tet methylcytosine dioxygenase genes *tet1*, *tet2*, and *tet3*, collectively known as the Tet enzymes, using CRISPR/Cas9. As the Tet enzymes convert 5mC to 5hmC, their knockout resulted in a marked reduction of 5hmC in the zebrafish brain, primarily in the midbrain and hindbrain regions, with the pineal gland (PG), torus longitudinalis (TL), RF(Arg-Phe)amide family 26 amino acid peptide (QRFP) neurons, glycine transporter 2 (glyt2) neuronal stripe 1 (GLYT2S1), and vesicular glutamate transporter 2 (vglut2) neuronal stripe 1 (VGLUT2S1) consistently downregulated in all or two out of the three knockouts (Figures 1H–1J).

We then verified whether EpiBrain-DNA can detect changes in 5mC. Exposure to decitabine, a DNA methyltransferase (DNMT) inhibitor, led to a global reduction of 5mC across the brain as expected (Figure 1K). Interestingly, the strongest reductions in 5mC were observed in brain regions that overlapped with those regulated by the Tet knockouts, particularly the pineal gland (PG), torus longitudinalis (TL), and GLYT2S1 and VGLUT2S1 neuronal clusters (Figures 1H-1K).

Counterstaining histone H3 with N-terminal histone H3 modifications on lysine 4 and 9, such as lysine 9 dimethylation (H3K9me2), was initially unsuccessful (Supplementary Figure 6A). We found that the anti-H3 antibody used for EpiBrain-H3(K27/K36) (Proteintech, Cat#68345) targets N-terminal residues of histone H3 that overlap the K4 and K9 amino acid positions. While this N-terminal-targeting anti-H3 antibody (hereafter referred to as anti-H3Tail) did not interfere with the detection of K27 and K36 modifications, we reasoned that the failure to co-stain with anti-K4 and anti-K9 antibodies was likely due to epitope competition between anti-H3Tail and antibodies recognizing the K4 and K9 modifications.

We therefore hypothesized that an alternative anti-H3 antibody targeting the C-terminal (core) region of histone H3 (hereafter referred to as anti-H3Core) would permit co-staining with anti-K4/K9 antibodies (Supplementary Figure 6B). Proper staining of the histone core required HCl-based denaturation, following the same protocol established for EpiBrain-DNA (Figure 1B). Indeed, an anti-H3Core antibody (Cell Signaling Technology, Cat#14269) targeting the C-terminal region of H3 successfully co-stained with antibodies against H3K9me2 (Supplementary Figure 7A), H3K9ac (Supplementary Figure 7B), H3K4me1 (Supplementary Figure 7C), and H3K4me3 (Supplementary Figure 7D). An H3Core template brain was generated by counterstaining wild-type larvae with the anti-H3Core antibody and anti-tErk antibody (Supplementary Figure 7E) and registering the H3Core channel to the tErk reference brain (Supplementary Figure 7F). We hereafter refer to this protocol as EpiBrain-H3(K4/K9) (Figure 1B).

To verify the effectiveness of the EpiBrain-H3(K4/K9) protocol, we first knocked out the zebrafish *setd7* gene (SET domain containing 7), which encodes a histone lysine methyltransferase that catalyzes methylation of histone H3 at lysine 4 (H3K4). As expected, *setd7* knockout led to reduced levels of lysine 4 monomethylation (H3K4me1), most prominently in the hindbrain gad1b neuronal cluster 14 (GAD1BC14) and VMAT2S1 (Figure 1L). In addition, lysine 4 trimethylation (H3K4me3) was reduced in both the forebrain and hindbrain, with particularly strong effects in forebrain subpallial dopaminergic clusters (SPDC) and in hindbrain tyrosine hydroxylase-positive neurons (TH) and noradrenergic neurons of the interfascicular and vagal areas (NA) (Figure 1M).

We next assessed whether changes in histone H3 lysine 9 modifications could be detected using EpiBrain-H3(K4/K9). Larvae were first exposed to UNC0642, a selective inhibitor of the lysine methyltransferases G9a (EHMT2) and GLP (EHMT1) with high potency in reducing the H3K9me2 mark. Consistent with its known activity, we detected a reduction in H3K9me2, most prominently in the pineal gland (PG) and torus longitudinalis (TL) (Figure 1N). To further validate the ability of the EpiBrain-H3(K4/K9) protocol to detect histone acetylation changes, we exposed larvae to C646, a selective inhibitor of the histone acetyltransferase p300. As expected, reduced lysine 9 acetylation (H3K9ac) was observed across the brain, with particularly pronounced reductions in the forebrain olfactory bulb (OB) and in hindbrain cerebellar gad1b-enriched areas (CGEA) and vglut2 neuronal stripe 2 (VGLUT2S2) (Figure 1O). Together, these results demonstrate that EpiBrain can reliably detect anticipated brain-wide epigenetic changes induced by perturbations with well-characterized epigenetic modulatory activities, often revealing previously unknown regional specificity in the process.

### EpiBrain detects changes in brain’s epigenetic landscape following chemical, genetic, and behavioral perturbations

Following the establishment of the EpiBrain method, we examined how the brain’s epigenetic landscape is modulated by factors relevant to human mental health and disease, including environmental toxicants, drugs of addiction, genetic mutations, and behavioral experiences. We first tested the effects of several prevalent environmental toxicants. We recently identified a new class of toxicants, topoisomerase II alpha (Top2a) inhibitors, that contribute to autism risk by modulating the H3K27me3 pathway^10^. Consistent with this mechanism, the Top2a inhibitor flumequine selectively altered H3K27me3 in the hindbrain, including QRFP neurons, VGLUT2S1, cerebellum gad1b-enriched area (CGAD1B), and hypocretin (orexin) receptor neurons HCRTRS4 (Figure 2A). Notably, exposure to the environmental toxicant bisphenol A (BPA), which has also been reported to exhibit Top2a inhibitory activity^41^, produced a highly similar pattern of H3K27me3 alteration in these same brain regions (Figure 2B). Among these, QRFP neurons and hypocretin receptor neurons were most strongly affected by BPA. Both QRFP and hypocretin signaling regulate appetite, and BPA exposure has been shown to affect appetite and potentially contribute to weight gain in humans^42,43^, rodents^44–46^, and zebrafish^47^, suggesting a potential H3K27me3-related epigenetic mechanism underlying this phenomenon. Flumequine and BPA treatments also induced broadly similar patterns of increased basal brain activity in the forebrain and midbrain, as revealed by pErk/tErk staining (Figures 2C & 2D), but with distinct regional biases. Specifically, subpallial otpb neuronal cluster 2 (OTPBC2) and midbrain hypocretin receptor neurons (MHCRTR) were most strongly activated following flumequine treatment (Figure 2C), whereas subpallial dopaminergic clusters (SPDC) and retinal arborization field 9 (AF9) showed the strongest activation following BPA exposure (Figure 2D). BPA has additionally been reported to inhibit DNA methylation in yeast^48^; in agreement with this observation, we detected a mild downregulation of DNA methylation (5mC and 5hmC) in scattered forebrain and midbrain regions, particularly in the habenula (HB), following BPA exposure (Figures 2E & 2F).

**Figure 2.**
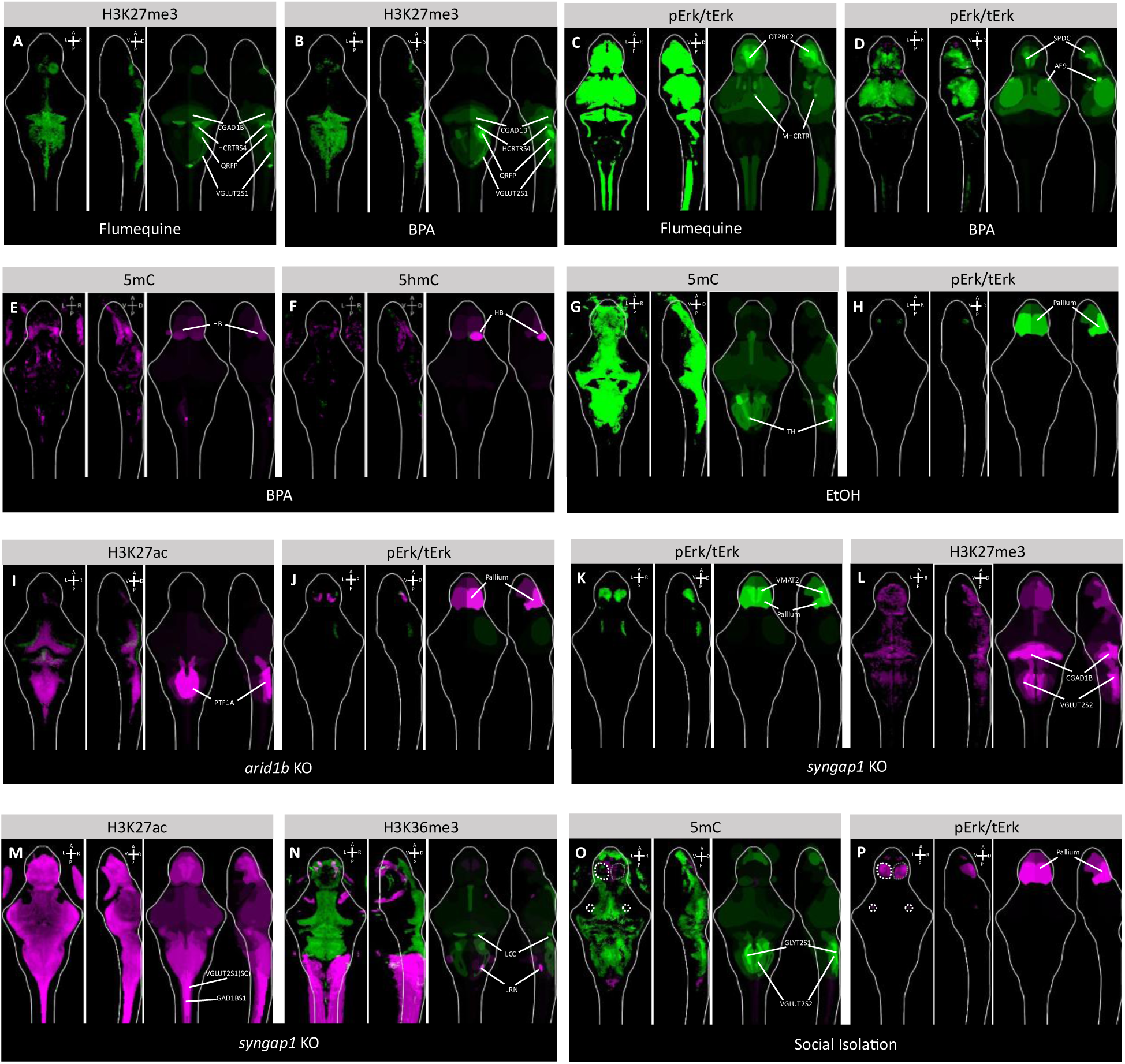
Chemicals, genetic mutations, and brain activities alter brain’s epigenetic landscape. (**A**) Embryonic exposure (0-3 dpf) to 25 µM Flumequine induced sustained upregulation of H3K27me3, mainly in the hindbrain, as detected by EpiBrain-H3(K27/K36). Two images on the left are dorsal and lateral views of the brain. Changes in the relative abundance of epigenetic modification are depicted by colors: green represents elevation compared to the control, and magenta represents reduction. A: anterior. P Posterior. L: left. R: right. V: ventral. D: dorsal. The image on the right highlights brain regions (dorsal and lateral views) with significant changes in epigenetic modification in colors. Green represents regions with elevated levels of epigenetic modification compared to the control, while magenta represents reduced levels of modification. Brightness of the colors depict the degree of significance. (**B**) Embryonic exposure (0-3 dpf) to 20 µM BPA induced upregulation of H3K27me3 in a pattern similar to the change following Flumequine treatment (A), as detected by EpiBrain-H3(K27/K36). (**C-D**) Embryonic exposures to Flumequine and BPA induced lasting elevation of basal brain activity in 6 dpf larvae, as detected by zBrain analysis through pErk/tErk staining. Activity increases are mainly restricted to the forebrain and midbrain regions. (**E-F**) Embryonic exposure (0-3 dpf) to 20 µM BPA downregulated the levels of 5mC (E) and 5hmC (F), primarily in the forebrain, as detected using the EpiBrain-DNA protocol. (**G**) A brief 2-hour embryonic exposure to 1% ethanol at 24 hpf induced a lasting upregulation of 5mC throughout the larval brain, as detected by EpiBrain-DNA. (**H**) Ethanol exposure upregulated basal brain activities of the pallium in the forebrain, as shown by pErk/tErk staining. (**I-J**) F0 CRISPR/Cas9 knock out (KO) of the *arid1b* gene resulted in significant reduction of H3K27ac in the caudal midbrain and hindbrain regions (GLYT2), as detected by EpiBrain-H3(K27/K36) (I). pErk/tErk staining revealed a forebrain structure within the pallium with partial suppression and partial elevation of basal brain activity (J). (**K-N**) F0 CRISPR/Cas9 double-KO of *syngap1a* and *syngap1b* genes (*syngap1*) led to upregulated basal brain activities in the forebrain as detected by pErk/tErk staining (K). EpiBrain-H3(K27/K36) detected brain-wide reductions in H3K27me3 (L) and H3K27ac (M), as well as elevated H3K26me3 in the brain and reduced H3K28me3 in the muscle (N). (**O-P**) Social isolation from 0-6 dpf led to pervasive elevation in 5mC as detected by EpiBrain-DNA (O). pErk/tErk staining detected reduced basal activity in parts of the forebrain and rostral lateral midbrain (P). White dashed circles enclose brain regions with reduced 5mC (O) as well as basal activity (P). Brain regions are identified based on alignment with the zBrain atlas following brain registration. The top differentially regulated brain regions are labeled in the images. Raw analysis results for all differentially regulated brain regions and their signal intensities in images shown in Figure 2 are summarized in Supplementary Data 2. AF9: retinal arborization field 9. HB: Habenula. CGAD1B: cerebellum gad1b enriched area. GAD1BS1: gad1b stripe 1 (spinal cord). GLYT2S1: glyt2 stripe 1. HCRTRS4: 6.7FDhcrtR-Gal4 stripe 4. LCC: lobus caudalis cerebeli. LRN: lateral reticular nucleus. MHCRTR: midbrain sparse 6.7FRhcrtR cluster. OTPBC2: subpallial Otpb cluster 2. PTF1A: ptf1a stripe. QRFP: qrfp neuron cluster sparse. SPDC: subpallial dopaminergic cluster. TH: small cluster of TH-stained neurons. TN: tectum neuropil. VGLUT2S1(SC): vglut2 stripe 1 (spinal cord). VGLUT2S2: vglut2 stripe 2 (rhombencephalon).

Chronic alcohol consumption is associated with alterations in brain DNA methylation^49^. Specifically, global increases in DNA methylation have been reported in patients with alcoholism^50^. In addition, fetal alcohol syndrome (FAS), which results from prenatal alcohol exposure and is characterized by neurodevelopmental impairments, has been linked to a distinct DNA methylation signature in affected individuals, with hypermethylated CpG sites enriched in genes regulating neurodevelopmental processes and associated with neurodevelopmental disorders^51^. To examine whether similar DNA methylation changes occur in a zebrafish model of FAS^52^, we exposed embryos at 24 hpf to 1% ethanol for two hours to model prenatal alcohol exposure. Ethanol-treated larvae exhibited a brain-wide increase in DNA methylation, with particularly pronounced effects in hindbrain tyrosine hydroxylase-positive neurons (TH), as detected by 5mC staining (Figure 2G). Prenatal ethanol exposure has been shown to significantly reduce TH⁺ dopamine-producing neurons in mouse^53^ and rat^54^, and our findings suggest a potential epigenetic contribution to this phenomenon through elevated 5mC levels. No significant changes in 5hmC were detected under these conditions (not shown), whereas elevated basal brain activity was observed in the forebrain pallium region (Figure 2H).

Our previous findings suggest that epigenetic dysregulation, particularly involving the H3K27me3 and histone acetylation pathways, are strongly associated with autism risk^9,10^. We therefore knocked out the zebrafish orthologs of two high-confidence autism risk genes, with or without known functional links to epigenetic regulation, to examine their effects on brain epigenetic states. Knocking out the zebrafish *arid1b* gene (orthologous to human *ARID1B*), a component of the SWI/SNF chromatin remodeling complex, resulted in a significant reduction of H3K27ac, as expected, most prominently in the caudal midbrain and hindbrain pancreas transcription factor 1α (ptf1a) neurons (PTF1A) (Figure 2I). This epigenetic change was accompanied by reduced basal brain activity in the forebrain pallium region (Figure 2J).

Interestingly, knockout of *syngap1a* and *syngap1b* (hereafter collectively referred to as *syngap1*), the zebrafish orthologs of the human *SYNGAP1* gene encoding a synaptic protein, not only altered basal brain activity in the forebrain pallium region and VMAT2 neuronal clusters (Figure 2K), but also induced widespread changes in multiple histone modifications, including H3K27me3 (Figure 2L), H3K27ac (Figure 2M), and H3K36me3 (Figure 2N). Specifically, H3K27me3 was reduced throughout the brain, with the strongest decreases observed in the CGAD1B and hindbrain VGLUT2S2 neurons (Figure 2L). H3K27ac was broadly reduced across the brain, most prominently in spinal cord vglut2 stripe 1 (VGLUT2S1(SC)) and spinal cord gad1b stripe 1 (GAD1BS1) neuronal populations (Figure 2M). In contrast, H3K36me3 was elevated in the midbrain and hindbrain, particularly in LCC neurons, and reduced in muscle tissue as well as in the hindbrain lateral reticular nucleus (LRN) (Figure 2N). As SYNGAP1 has no known direct role in epigenetic regulation, these findings suggest that alterations in synaptic function and neuronal activity can secondarily reshape the brain’s epigenetic landscape, a hypothesis that we further explore in the following sections.

In addition to chemical and genetic perturbations, early life experiences have also been associated with long-lasting changes in the brain’s epigenetic state. For example, social isolation has been shown to alter DNA methylation in the mouse brain^55^. To determine whether similar effects occur in zebrafish, we socially isolated larvae from 0 to 6 dpf by raising each individual in a separate well of an opaque 96-well plate. This manipulation resulted in a widespread increase in 5mC across the brain, with the exception of the medial forebrain, rostral lateral midbrain, and rostral spinal cord regions (Figure 2O). The increase in 5mC was particularly pronounced in the hindbrain, especially within GLYT2S1 and VGLUT2S2 neuronal clusters (Figure 2O). No significant changes in 5hmC levels were detected under these conditions (not shown). Notably, reduced basal brain activity was observed in the medial forebrain pallium regions and rostral midbrain (Figure 2P; regions enclosed by dashed circles), which partially overlapped with the brain regions that did not exhibit increased 5mC following social isolation (Figure 2O; regions enclosed by dashed circles).

### Changes in brain activity induce lasting changes in brain’s epigenetic landscape

While chemicals and genetic perturbations with known epigenetic modulatory activities are expected to directly influence the brain’s epigenetic state, the mechanisms by which synaptic proteins or behavioral experiences alter epigenetic pathways are less clear (Figures 2L–2O). Similarly, it remains poorly understood whether drugs of addiction modify brain epigenetics primarily through direct chemical interactions with epigenetic machinery or indirectly via their psychoactive effects^56–58^. We therefore hypothesized that changes in brain activity itself can, at least in part, mediate the epigenetic alterations induced by these activity-modulating perturbations.

We tested this hypothesis by inducing brain-wide neuronal activation using the chemical convulsant pentylenetetrazol (PTZ)^59^ and examining its impact on the brain’s epigenetic state using EpiBrain. We first employed pErk/tErk staining to confirm the effectiveness of PTZ in eliciting seizure-like neuronal activity and found that a 1-hour 15 mM PTZ treatment at 6 dpf induced widespread brain activation, particularly in the forebrain pallium and midbrain tectum neuropil (TN) regions (Figure 3A). Notably, a sustained elevation in baseline brain activity was detected 24 hours after a 1-hour PTZ exposure at 5 dpf (Figure 3B). This persistent activity increase was again most prominent in the forebrain (most prominently in the VMAT2 and SPDC neurons) and midbrain tectum neuropil (Figure 3B) and substantially overlapped with the activity pattern observed immediately after PTZ treatment (Figure 3A), indicating that a brief, transient episode of seizure-like activity is sufficient to induce lasting changes in brain activity.

**Figure 3.**
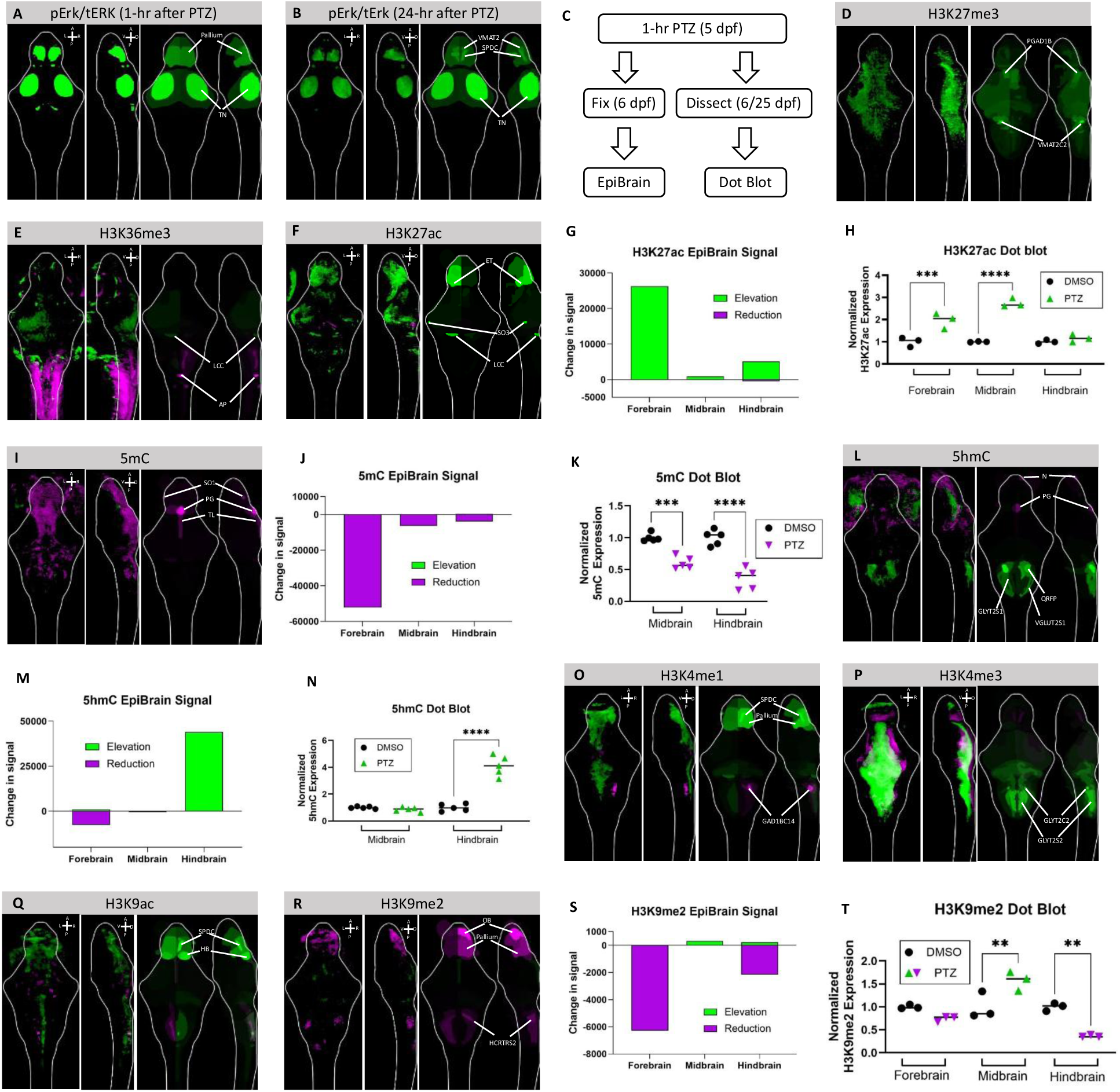
PTZ induces brain-wide changes in epigenetic modifications. (**A-B**) pErk/tErk staining detected brain-wide activity immediately after 1-hour PTZ treatment (A) or 24-hours after the 1-hour PTZ exposure (B). Green denotes brain regions that are hyperactivated compared to the DMSO-treated control. Magenta denotes brain regions that are hypoactivated compared to the control. (**C**) A schematic illustration of experimental design. On 6 dpf, 24 hours following a 1-hour PTZ treatment on 5 dpf, zebrafish larvae are rapidly fixed for EpiBrain staining. On 6 or 25 dpf, 24 hours following a 1-hour PTZ treatment on 5 or 24 dpf, zebrafish larvae are dissected for dot blot analysis. (**D-F**) EpiBrain-H3(K27/K36) staining and imaging analysis detected lasting changes in the levels of H3K27me3 (D), H3K36me3 (E), and H3K27ac (F) 24-hours post-PTZ treatment. Brain regions were labeled based on alignment with the zBrain atlas following brain registration. The two images on the left are dorsal and lateral views of the brain. Changes in the relative abundance of epigenetic modification are depicted by colors: green represents elevation compared to the control, and magenta represents reduction. A: anterior. P Posterior. L: left. R: right. V: ventral. D: dorsal. The image on the right highlights brain regions with significant changes in epigenetic modification in colors, in dorsal and lateral views. Green represents regions with elevated levels of epigenetic modification compared to the control; magenta represents reduced levels of modification. Brightness of the colors depict the degree of significance. (**G**) Bar plot showing the summed changes in H3K27ac signal across the forebrain, midbrain, and hindbrain following PTZ exposure. For each major brain subdivision, all regions of interest exhibiting statistically significant H3K27ac changes were identified, and the EpiBrain signal changes within each subdivision were summed to generate a single aggregate value per region. Positive values (green) indicate net elevation of H3K27ac, whereas negative values (magenta) indicate net reduction. This analysis reveals pronounced H3K27ac elevation in the forebrain, with comparatively modest changes in the midbrain and hindbrain. (**H**) Dot blot analysis of brain samples collected from 25 dpf larvae pre-exposed to PTZ for 1-hour at 24 dpf detected significantly elevated H3K27ac in the forebrain and midbrain. No significant changes in H3K27ac were detected in the hindbrain. Green: elevation. Magenta: reduction. Significance was calculated by one-way ANOVA and Tukey’s multiple comparisons test. \*\*\**P* < 0.001; \*\*\*\**P* < 0.0001. (**I**) EpiBrain-DNA detected lasting changes in the level of 5mC 24-hours post-PTZ treatment. Green: elevation. Magenta: reduction. (**J**) Bar plot showing the summed changes in 5mC signal across the forebrain, midbrain, and hindbrain following PTZ exposure. For each major brain subdivision, all regions of interest exhibiting statistically significant 5mC changes were identified, and the EpiBrain signal changes within each subdivision were summed to generate a single aggregate value per region. Positive values (green) indicate net elevation of 5mC, whereas negative values (magenta) indicate net reduction. This analysis reveals a pronounced reduction in 5mC in the forebrain, with more modest decreases in the midbrain and hindbrain. (**K**) Dot blot analysis of brain samples collected from 6 dpf larvae pre-exposed to PTZ for 1-hour at 5 dpf detected significant reductions of 5mC in the midbrain and hindbrain regions. Green: elevation. Magenta: reduction. Significance was calculated by one-way ANOVA and Tukey’s multiple comparisons test. \*\*\**P* < 0.001; \*\*\*\**P* < 0.0001. (**L**) EpiBrain-DNA detected lasting changes in the level of 5hmC 24-hours post-PTZ treatment. Green: elevation. Magenta: reduction. (**M**) Bar plot showing the summed changes in 5hmC signal across the forebrain, midbrain, and hindbrain following PTZ exposure. For each major brain subdivision, all regions of interest exhibiting statistically significant 5hmC changes were identified, and the EpiBrain signal changes within each subdivision were summed to generate a single aggregate value per region. Positive values (green) indicate net elevation of 5hmC, whereas negative values (magenta) indicate net reduction. This analysis reveals a pronounced increase in 5hmC in the hindbrain, accompanied by a modest reduction in the forebrain and minimal net change in the midbrain. (**N**) Dot blot analysis of brain samples collected from 6 dpf larvae pre-exposed to PTZ for 1-hour at 5 dpf detected significant elevation of 5mC in the hindbrain, but not the forebrain and midbrain. Green: elevation. Magenta: reduction. Significance was calculated by one-way ANOVA and Tukey’s multiple comparisons test. \*\*\*\**P* < 0.0001. (**O-R**) EpiBrain-H3(K4/K9) staining and image analysis detected lasting changes in the levels of H3K4me1 (O), H3K4me3 (P), H3K9ac (Q), and H3K9me2 (R) 24-hours post-PTZ treatment. Green: elevation. Magenta: reduction. (**S**) Bar plot showing the summed changes in H3K9me2 signal across the forebrain, midbrain, and hindbrain following PTZ exposure. For each major brain subdivision, all regions of interest exhibiting statistically significant H3K9me2 changes were identified, and the EpiBrain signal changes within each subdivision were summed to generate a single aggregate value per region. Positive values (green) indicate net elevation of H3K9me2, whereas negative values (magenta) indicate net reduction. This analysis reveals a strong reduction of H3K9me2 in the forebrain and hindbrain, accompanied by a modest increase in the midbrain. (**T**) Dot blot analysis of brain samples collected from 25 dpf larvae pre-exposed to PTZ for 1-hour at 24 dpf detected a significant elevation of H3K9me2 in the midbrain and a significant reduction of H3K9me2 in the hindbrain. Green: elevation. Magenta: reduction. Significance was calculated by one-way ANOVA and Tukey’s multiple comparisons test. \*\**P* < 0.01. Brain regions are identified based on alignment with the zBrain atlas following brain registration. The top differentially regulated brain regions are labeled in the images. Raw analysis results for all differentially regulated brain regions and their signal intensities in images shown in Figure 3 are summarized in Supplementary Data 3. ET: eminentia thalami. GAD1BC14: gad1b cluster 14. GLYT2C2: glyt2 cluster 2. GLYT2S1: glyt2 stripe 1. GLYT2S2: glyt2 stripe 2. HCRTRS2: 6.7FDhcrtR-Gal4 stripe 2. N: lateral line neuromast N ganglia. PG: pineal gland. PGAD1B: pretectal gad1b cluster. QRFP: qrfp neuron cluster sparse. SO1: lateral line neuromast so1 ganglia. SO3: lateral line neuromast so3 ganglia. SPDC: subpallial dopaminergic cluster. TL: torus longitudinalis. VGLUT2S1: vglut2 stripe 1. VMAT2C2: vmat2 cluster 2 (rhombencephalon).

To determine whether PTZ treatment induces epigenetic changes, larval zebrafish were collected 24 hours post-PTZ exposure for EpiBrain analysis and independent validation using dot blot assays (Figure 3C). Significant alterations were detected across all epigenetic markers examined using the three EpiBrain protocols (Figure 1B). Using the EpiBrain-H3(K27/K36) protocol, we observed a brain-wide increase in H3K27me3, most prominently in the pretectal gad1b cluster (PGAD1B) and the hindbrain vmat2 neuronal cluster 2 (VMAT2C2) (Figure 3D). Regionally patterned increases and decreases were observed for H3K36me3, including an increase in LCC and a decrease in AP in the brain, as well as decreases in adjacent muscle tissues (Figure 3E). Sporadic increases in H3K27ac were detected in the forebrain eminentia thalami (ET), lateral line neuromast so3 ganglia (SO3), and hindbrain LCC regions (Figure 3F). Quantification of the H3K27ac EpiBrain images revealed that H3K27ac elevation was primarily localized to the forebrain (Figure 3G). To validate this result, brains from 25 dpf zebrafish that had been subjected to a 1-hour PTZ treatment at 24 dpf were microdissected and separated into forebrain, midbrain, and hindbrain fragments. Zebrafish at 25 dpf were used to ensure sufficient tissue yield and to assess conservation of the observed epigenetic responses at a later developmental stage. Dot blot analysis confirmed that H3K27ac levels were significantly increased in the forebrain, as expected based on the EpiBrain result (Figure 3G), and also revealed an unexpected but significant increase in the midbrain, with no significant changes detected in the hindbrain (Figure 3H).

Using the EpiBrain-DNA protocol, we detected a pervasive reduction in 5mC across the brain (Figures 3I & 3J), with particularly strong decreases in the lateral line neuromast so1 ganglia (SO1) and forebrain pineal gland (PG) and torus longitudinalis (TL) regions but also scattered reductions in midbrain and hindbrain (Figure 3I). These findings were corroborated by dot blot analysis of midbrain and hindbrain samples microdissected from 6 dpf larvae following a 1-hour PTZ treatment at 5 dpf, which revealed significantly reduced 5mC levels in both the midbrain and hindbrain (Figure 3K). In addition, EpiBrain-DNA revealed a significant increase in 5hmC in the hindbrain QRFP, GLYT2S1, and VGLUT2S1 neurons as well as modest reductions in the lateral line neuromast N ganglia (N) and the forebrain pineal gland (Figures 3L & 3M), findings that were likewise confirmed by dot blot analysis (Figure 3N).

Finally, the EpiBrain-H3(K4/K9) protocol detected region-specific changes in H3K4me1 (Figure 3O), H3K4me3 (Figure 3P), H3K9ac (Figure 3Q), and H3K9me2 (Figure 3R). Specifically, EpiBrain revealed increased H3K4me1 in the forebrain SPDC and pallium regions and decreased H3K4me1 in the hindbrain GAD1BC14 (Figure 3O). Increased H3K4me3 was observed in the hindbrain glyt2 cluster 2 (GLYT2C2) and glyt2 stripe 2 (GLYT2S2) neuronal clusters (Figure 3P). H3K9ac was found to be increased in the forebrain SPDC and habenula (Figure 3Q). Finally, EpiBrain revealed reduced H3K9me2 levels in the forebrain olfactory bulb and pallium as well as hindbrain 6.7FDhcrtR-Gal4 stripe 2 (HCRTRS2) neuronal clusters, accompanied by sparsely elevated H3K9me2 in the midbrain (Figures 3R & 3S). This pattern was largely confirmed by dot blot analysis of brain samples collected from 25 dpf larvae following a 1-hour PTZ treatment at 24 dpf, which showed reduced H3K9me2 in forebrain and hindbrain tissues and elevated H3K9me2 in the midbrain (Figure 3T). Together, these results demonstrate that a brief episode of seizure-like neuronal activity is sufficient to induce persistent, brain-region-specific alterations across multiple histone and DNA epigenetic marks, supporting our hypothesis that changes in brain activity itself can drive lasting remodeling of the brain’s epigenetic landscape.

### The epigenetic-modulatory immediate early gene *gadd45b* mediates activity-induced changes in 5hmC

Immediate early genes (IEGs) are rapidly induced in response to neuronal activation. A subset of IEGs have established roles in epigenetic regulation^60–62^, which we hereafter refer to as epigenetic immediate early genes (Epi-IEGs), providing a potential mechanistic link between neuronal activity and epigenetic remodeling. We therefore hypothesized that Epi-IEGs mediate activity-induced epigenetic changes in the brain.

To test this hypothesis, we induced brain-wide seizure-like activity in 6 dpf zebrafish larvae using PTZ and collected mRNA samples immediately following a 1-hour PTZ exposure to search for Epi-IEGs induced by seizure-like neuronal activity. RNA sequencing (RNA-seq) identified well-characterized IEGs as expected, including *fosab*, the zebrafish ortholog of the *Fos* gene. Among the 398 significantly upregulated and downregulated candidate IEGs detected by RNA-seq, several have reported roles in epigenetic regulation and were therefore classified as Epi-IEGs. Notably, *gadd45ba* and *gadd45bb* (hereafter collectively referred to as *gadd45b*) are the two zebrafish orthologs of *Gadd45b*, an Epi-IEG known to mediate the conversion of 5mC to 5hmC^61^ (Figure 4A). Additional Epi-IEG candidates included zebrafish orthologs of the mammalian genes *Gadd45a*^63^, *Prdm13*^64^, *Npas4*^65^, *Egr1*^66–68^, *Egr2*^69^, and *Jdp2*^70^ (Figure 4A). Quantitative PCR (qPCR) confirmed robust induction of *gadd45b* following PTZ treatment in dissected brain tissue (Figure 4B). Hybridization chain reaction (HCR)^71^ further revealed widespread expression of *gadd45ba* (Supplementary Figure 8A) and *gadd45bb* (*not shown*) throughout the 6 dpf larval brain. By co-staining *gadd45b* with *elavl3* (Supplementary Figure 8B) and registering the images to an *elavl3* reference brain (Supplementary Figure 8C), we mapped activity-dependent changes in *gadd45b* expression using the zBrain atlas and observed elevated *gadd45ba and gadd45bb* mRNA levels across the brain following PTZ treatment (Figure 4C).

**Figure 4.**
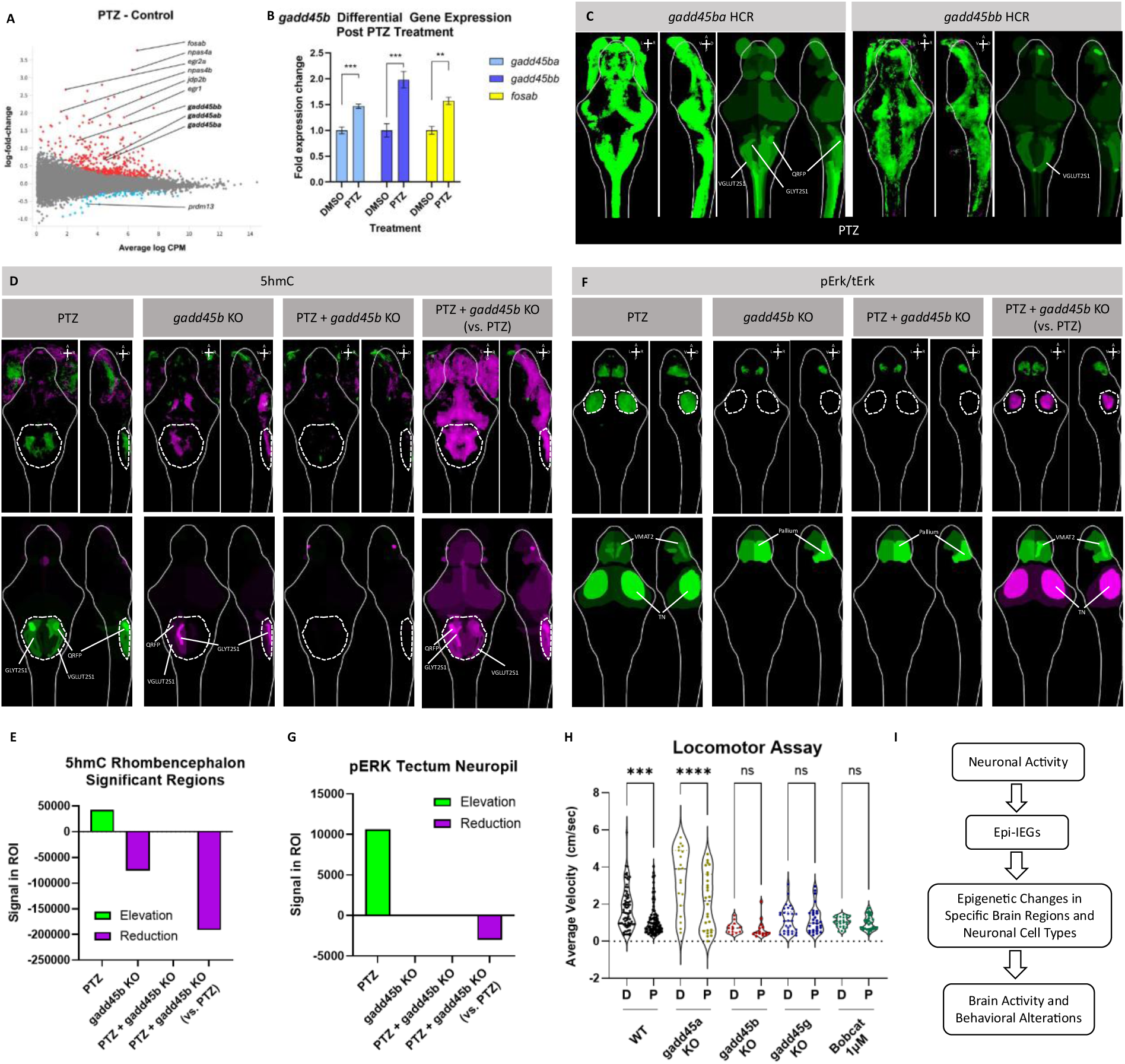
*gadd45b* mediates activity-induced changes in 5hmC. (**A**) RNA-sequencing detected 398 immediate early genes with significant changes in expression 1-hour post-PTZ treatment, including the following genes with reported epigenetic modulatory functions: *egr1*, *egr2a*, *gadd45ab*, *gadd45ba*, *gadd45bb*, *npas4a*, *npas4b*, and *prdm13*. The *fosab* (the zebrafish ortholog of *Fos*) gene was highly expressed as expected. (**B**) Quantitative PCR detected elevated expression of *gadd45ba* and *gadd45bb* in the 6 dpf larval zebrafish brain 1-hour following PTZ treatment, verifying the RNA-sequencing result. *fosab* expression was also significantly elevated and is shown as a positive control. Significance was calculated by two-tailed Student’s *t* test. \*\**P* < 0.01; \*\*\**P* < 0.001. (**C**) Hybridization chain reaction (HCR) detected brain-wide expressions of *gadd45ba* and gadd45bb in the 6 dpf larval zebrafish brain 1-hour following PTZ treatment. The increase in expressions were most significantly in the hindbrain regions. The HCR images were counter-stained for *elevl3* and registered to an *elaval3* reference brain from the zBrain database. Brain regions were labeled based on alignment with the zBrain atlas following brain registration. The two images on the left are dorsal and lateral views of the brain. Changes in the relative abundance of gene expression are depicted by colors: green represents elevation compared to the control, and magenta represents reduction. A: anterior. P Posterior. L: left. R: right. V: ventral. D: dorsal. The image on the right highlights brain regions with significant changes in gene expression in colors, shown in dorsal and lateral views. Green represents regions with elevated levels of gene expression compared to the control; magenta represents reduced levels of gene expression. Brightness of the colors depict the degree of significance. (**D**) EpiBrain-H3(K4/K9) imaging and analysis detected changes in 5hmC, especially in the hindbrain regions (white dashed circles), following various treatment conditions. Specifically, PTZ-induced an elevation of 5hmC in the hindbrain, which was rescued by *gadd45b* KO. Green: elevation. Magenta: reduction. PTZ: 1-hr PTZ treatment at 5 dpf; analysis was conducted by comparing to DMSO control; re-adaptation of Figure 3L. *gadd45b* KO: F0 CRISPR/Cas9 double-KO of *gadd45ba* and *gadd45bb*, analysis was conducted by comparing to wild-type (WT) control. PTZ + *gadd45b* KO: *gadd45b* KO larvae exposed to PTZ for 1-hr at 5 dpf; analysis was conducted by comparing to WT control. PTZ + *gadd45b* KO (vs. PTZ): *gadd45b* KO larvae exposed to PTZ for 1-hr at 5 dpf; analysis was conducted by comparing to WT control exposed to PTZ for 1-hr at 5 dpf. (**E**) Bar plot showing summed 5hmC signal changes in rhombencephalon regions of interest (ROIs; white dashed circles in Figure 4D) following PTZ exposure, *gadd45b* KO, combined PTZ + *gadd45b* KO, and PTZ + *gadd45b* KO relative to PTZ alone. Rhombencephalon regions were first identified based on significant 5hmC changes in each condition, and only regions consistently detected across all comparisons were included in the analysis. For each condition, 5hmC signal was summed across these shared ROIs and plotted, with positive values indicating elevation and negative values indicating reduction in 5hmC signal. The analysis reveals PTZ-induced increases in 5hmC (PTZ) that are reversed by *gadd45b* knockout (PTZ + *gadd45b* KO). (**F**) pErk/tErk staining and analysis revealed changes in basal brain activity, particularly in the midbrain optic tectum area (white dashed circles), following various treatment conditions. Specifically, PTZ-induced an increase in basal activity in the tectum, a change that was blocked by *gadd45b* KO. Green: elevation. Magenta: reduction. TN: tectum neuropil. PTZ: 1-hr PTZ treatment at 5 dpf; analysis was conducted by comparing to DMSO control; re-adaptation of Figure 3B. *gadd45b* KO: F0 CRISPR/Cas9 double-KO of *gadd45ba* and *gadd45bb*, analysis was conducted by comparing to wild-type (WT) control. PTZ + *gadd45b* KO: *gadd45b* KO larvae exposed to PTZ for 1-hr at 5 dpf; analysis was conducted by comparing to WT control. PTZ + *gadd45b* KO (vs. PTZ): *gadd45b* KO larvae exposed to PTZ for 1-hr at 5 dpf; analysis was conducted by comparing to WT control exposed to PTZ for 1-hr at 5 dpf. (**G**) Bar plot showing summed pERK signal changes in the tectum neuropil (white dashed circles in Figure 4F) following PTZ exposure, *gadd45b* KO, combined PTZ + *gadd45b* KO, and PTZ + *gadd45b* KO relative to PTZ alone. Positive values indicate elevation and negative values indicate reduction in pERK signal. The analysis reveals PTZ-induced increases in brain activity (pERK signal) (PTZ) that are reversed by *gadd45b* knockout (PTZ + *gadd45b* KO). (**H**) A 1-hour PTZ treatment at 5 dpf significantly reduced locomotor activity (average velocity) measured at 6 dpf. This PTZ-induced locomotor deficit was rescued by knockout of *gadd45b* or *gadd45g*, as indicated by the absence of significant differences between vehicle-treated (D: DMSO) and PTZ-treated groups (P: PTZ), whereas *gadd45a* knockout did not restore locomotor activity. In parallel, pharmacological inhibition of TET enzymes with Bobcat 339 (1 µM) via a 1-hour co-treatment during PTZ exposure similarly rescued the locomotor deficit. Significance was calculated by one-way ANOVA and Šídák’s multiple comparisons test. ns, not significant; \*\*\**P* < 0.001; \*\*\*\**P* < 0.0001. (**I**) A theoretical framework for how prior experience (in the form of neuronal activity) influence brain’s epigenetic state via IEGs with epigenetic modulatory function (Epi-IEGs), and subsequently influence brain activity and behavior. Brain regions are identified based on alignment with the zBrain atlas following brain registration. The top differentially regulated brain regions are labeled in the images. Raw analysis results for all differentially regulated brain regions and their signal intensities in images shown in Figure 4 are summarized in Supplementary Data 4. GLYT2S1: glyt2 stripe 1. QRFP: qrfp neuron cluster sparse. TN: tectum neuropil. VGLUT2S1: vglut2 stripe 1. VMAT2: vmat2 cluster (telencephalon).

To evaluate the role of *gadd45b* in activity-induced epigenetic changes, we generated *gadd45b* knockouts using CRISPR/Cas9. Given that mammalian *Gadd45b* has been implicated in DNA demethylation^61^, we assessed the impact of *gadd45b* loss on 5hmC levels using EpiBrain. Following PTZ exposure 24 hours prior to EpiBrain analysis (Figure 3C), a condition that elevates 5hmC in the hindbrain of wild-type fish, particularly in QRFP, GLYT2S1, and VGLUT2S1 neuronal populations (Figure 3L; shown again in Figure 4D: PTZ), *gadd45b* knockout completely abolished the PTZ-induced increase in 5hmC in these regions (Figures 4D & 4E: PTZ + *gadd45b* KO, regions enclosed by dashed circles). Loss of *gadd45b* alone also reduced baseline 5hmC levels in the same hindbrain regions (Figures 4D & 4E: *gadd45b* KO), an effect that was most pronounced when directly comparing PTZ-treated *gadd45b* knockout fish with PTZ-treated wild-type controls (Figures 4D & 4E: PTZ + *gadd45b* KO vs. PTZ).

In contrast, knockout of *gadd45a* (*gadd45aa* and *gadd45ab*) (Supplementary Figure 9A) or *gadd45g* (*gadd45ga* and *gadd45gb*) (Supplementary Figure 9B) reduced but did not fully eliminate the PTZ-induced elevation of 5hmC in the hindbrain (Supplementary Figures 9A & 9C: PTZ + *gadd45a* KO; Supplementary Figures 9B & 9D: PTZ + *gadd45g* KO). Notably, knockout of the methylcytosine dioxygenases *tet1* and *tet2* also led to depletion of 5hmC in similar hindbrain regions, particularly within the QRFP, GLYT2S1, and VGLUT2S1 neuronal clusters (Figures 1H & 1I).

We further hypothesized that the persistent changes in neuronal activity observed after PTZ exposure (Figure 3B; shown again in Figure 4F and Supplementary Figures 10A & 10B: PTZ) are driven, at least in part, by *gadd45b*-mediated, activity-induced epigenetic remodeling. Consistent with this idea, *gadd45b* knockout completely abolished the PTZ-induced increase in baseline activity within the tectum neuropil (TN) (Figures 4F & 4G: PTZ + *gadd45b* KO, regions enclosed by dashed circles). In contrast, *gadd45b* knockout did not effectively attenuate PTZ-induced activity in the forebrain (Figure 4F: PTZ + *gadd45b* KO); in fact, *gadd45b* knockout alone produced a modest increase in basal forebrain activity (Figure 4F: *gadd45b* KO), similar to the effect observed following PTZ treatment (Figure 4F: PTZ).

In comparison, knockout of *gadd45a* (Supplementary Figure 10A) or *gadd45g* (Supplementary Figure 10B) led to partial reductions in PTZ-induced activity within the tectum neuropil (Supplementary Figures 10A & 10C: PTZ + *gadd45a* KO; Supplementary Figures 10B & 10D: PTZ + *gadd45g* KO), but, unlike *gadd45b* knockout (Figures 4F & 4G: PTZ + *gadd45b* KO), did not fully abolish this activity. Similar to *gadd45b* knockout (Figure 4F), loss of *gadd45a* or *gadd45g* alone elicited increased basal forebrain activity (Supplementary Figure 10A: *gadd45a* KO; Supplementary Figure 10B: *gadd45g* KO) and failed to rescue PTZ-induced activity in this region (Supplementary Figure 10A: PTZ + *gadd45a* KO; Supplementary Figure 10B: PTZ + *gadd45g* KO).

We next examined whether the persistent changes in brain activity induced by PTZ exposure translate into behavioral alterations, and whether such effects can be rescued by inhibiting the *gadd45b*-*Tet*-5hmC pathway^61^. As described above, 5hmC levels in the QRFP neurons are selectively regulated by *gadd45b*, and the neuropeptide QRFP has been implicated in the control of locomotor activity and sleep in zebrafish^72^. Moreover, the PTZ-induced increase in basal activity within the tectum neuropil is selectively blocked by *gadd45b* knockout (Figures 4F & 4G: PTZ + *gadd45b* KO), and the optic tectum is a key structure regulating visuomotor behavior in zebrafish^73,74^. We therefore tested whether PTZ exposure induces lasting changes in locomotor behavior and whether these changes can be rescued genetically or pharmacologically.

Consistent with this hypothesis, larvae exposed to a 1-hour PTZ treatment at 5 dpf exhibited reduced locomotor activity at 6 dpf. This locomotor deficit was rescued by knockout of *gadd45b* as well as *gadd45g*, as shown by lack of significant differences in locomotor activities between control and PTZ-treated groups, whereas knockout of *gadd45a* did not restore PTZ-inhibited locomotor activity compared to the vehicle control (Figure 4H). In parallel, pharmacological inhibition of TET enzymes using Bobcat 339 similarly rescued the PTZ-induced locomotor deficit (Figure 4H). Together, these findings demonstrate that prior neuronal activity, such as seizure-like activity elicited by PTZ, can drive persistent changes in the brain’s epigenetic state, neural activity, and behavior through the activity of Epi-IEGs such as *gadd45b* (Figure 4I).

## DISCUSSION

In this study, we introduce EpiBrain, a whole-brain imaging and registration platform that enables the visualization of epigenetic changes across the zebrafish brain at cellular resolution. By integrating immunohistochemistry, volumetric imaging, and brain atlas-based registration, EpiBrain provides an unbiased snapshot of the brain’s epigenetic landscape in response to diverse physiological and pathological perturbations. Using this approach, we uncovered brain-wide epigenetic changes induced by chemical exposures, genetic mutations, and neuronal activity, and identified the immediate early gene *gadd45b* as an Epi-IEG and a key mediator of activity-dependent DNA demethylation and behavioral modulation.

A key innovation of EpiBrain is its ability to resolve brain region-specific changes in histone and DNA modifications in intact larval zebrafish brains. Traditional epigenetic profiling techniques, such as ChIP-seq, CUT&RUN, bisulfite sequencing, and bulk biochemical assays, require tissue homogenization prior to the assay and are unable to detect the spatial distribution of epigenetic changes. By contrast, EpiBrain retains the anatomical context of epigenetic changes, allowing researchers to pinpoint specific neuronal populations or brain regions affected by experimental manipulations. Our results demonstrate the utility of EpiBrain in detecting both expected and novel patterns of epigenetic modulation. For instance, we verified known drug-induced changes in histone modifications using inhibitors targeting EZH2 (Figure 1C), KDM6A/B (Figure 1D), p300 (Figure 1E) and HDACs (Figure 1F), while also uncovering unexpected region-specific effects of environmental toxicants, such as BPA and flumequine, on H3K27me3 in hindbrain appetite-regulating neurons (Figures 2A & 2B).

Our findings also establish EpiBrain as a valuable tool for mechanistic studies. By combining EpiBrain with genetic perturbation and behavioral assays, we show that seizure-like activity induced by PTZ elicits durable changes in multiple epigenetic marks, including histone and DNA modifications, in distinct brain regions (Figures 3D-3T). Importantly, we demonstrate that *gadd45b*, a brain-expressed epigenetic immediate early gene (Epi-IEG), is required to mediate PTZ-induced increases in 5hmC in the hindbrain. Loss of *gadd45b* not only blocked 5hmC elevation (Figures 4D & 4E) but also reversed sustained increases in tectal neural activity (Figures 4F & 4G) and rescued behavioral deficits in locomotion. These findings provide a direct mechanistic link between neuronal activity, epigenetic remodeling, and behavioral outcomes, demonstrating the importance of Epi-IEGs in translating environmental and physiological stimuli into long-lasting epigenetic changes.

EpiBrain also sheds light on how synaptic genes not traditionally associated with epigenetic regulation, such as *SYNGAP1*, can nonetheless drive profound alterations in histone methylation and acetylation patterns. This supports a model in which neural activity and circuit function feedback to reshape the epigenetic state of the brain, potentially via activity-dependent transcriptional programs involving Epi-IEGs. The bidirectional interplay between synaptic signaling and chromatin remodeling revealed by our data provides a framework for understanding how genetic and environmental factors converge to influence neurodevelopmental trajectories and mental health.

While our current implementation of EpiBrain is limited to larval zebrafish, the method is broadly extensible. Similar imaging-based approaches could be adapted for use in mammalian systems, including rodents, where brain-wide epigenetic mapping remains technically challenging. Moreover, future iterations of EpiBrain may incorporate additional molecular modalities, such as multiplexed labeling or spatial transcriptomics, to deepen our understanding of the molecular consequences of epigenetic changes.

In conclusion, EpiBrain provides a rapid, robust, and scalable method for whole-brain analysis of epigenetic states *in vivo*. By enabling spatially resolved detection of epigenetic modifications in response to diverse perturbations, EpiBrain fills a critical gap in neuroepigenetics research and opens new avenues for investigating the molecular basis of brain plasticity, development, and disease.

## MATERIALS AND METHODS

### Zebrafish husbandry

Zebrafish were housed at 26°C–27°C on a 14-hour light, 10-hour dark light cycle. Wild-type AB strain was used for all experiments. All zebrafish experiments were approved by the Institutional Animal Care and Use Committee at the University of Washington.

### EpiBrain staining and imaging

Larvae at 6 dpf were fixed in cold (4°C) 4% paraformaldehyde (PFA) in phosphate buffered saline (PBS) containing 0.025% Triton-X-100 (PBT). For a typical experiment, n≥20 larvae were collected to ensure sufficient power for image analysis. The samples were incubated in the PFA solution overnight and then rinsed in PBT. PBT was used for all subsequent rinses.

For EpiBrain-H3(K27/K36), following rinsing from the PFA solution, the fish were bleached for 30 minutes in water containing 3% H_2_O_2_ and 1% KOH to remove pigmentation. After rinsing, the tissue underwent antigen retrieval in 150 mM Tris-HCl buffer (pH 9.0) for 15 minutes at 70°C. The fish were then permeabilized with TrypLE Express Enzyme Solution (Gibco) (diluted 1:5 in PBS) at 4°C, blocked in blocking buffer (PBT + 1% dimethyl sulfoxide (DMSO) + 1% bovine serum albumin (BSA)) supplemented with 2% donkey serum for 1 hour at 4°C, and subsequently incubated with primary and secondary antibodies for 3 nights each in blocking buffer containing 1% heptakis(2,6-di-O-methyl)-β-cyclodextrin (DIMEB) (Santa Cruz Biotechnologies, CAT# sc-215141). The Proteintech anti-H3 antibody (CAT#68345) was used to counterstain H3K27 and H3K36 modifications.

For EpiBrain-H3(K4/K9) and EpiBrain-DNA, after rinsing from PFA, the fish were incubated in acrylamide hydrogel solution for 24 hours at 4°C. Following incubation, polymerization was achieved by transferring the fish in hydrogel solution to a 37°C incubator for 3 hours. Samples were then incubated in 8% SDS in boric acid buffer at 37°C for 3 days. Post-clearing, the fish were fixed again in 4% PFA for 20 minutes and rinsed. The following day, samples were bleached for 30 minutes and incubated in 2N HCl for 30 minutes at 37°C. After thoroughly rinsing out the HCl, the tissue was blocked in blocking buffer with 2% donkey serum for 1 hour at 37°C, then incubated in blocking buffer with 1% DIMEB and the primary and secondary antibodies at the desired concentrations for 4 nights each at 37°C. To enhance antibody penetration, the fish were centrifuged in each antibody solution for 3 hours at 400 × g. The Cell Signaling Technology anti-H3 antibody (CST#14269) which targets the carboxy terminus (the core region) of H3 was used to counterstain H3K4 and H3K9 modifications. If counterstaining 5mC/5hmC with propidium iodide (PI), incubate in 0.2mg/ml PI in PBT overnight at 37°C, and rinse thoroughly before imaging.

Fish were mounted dorsal side down in a 2% agarose gel mold and covered in 1% agarose before imaging by following the zMold mounting procedure^75^. Imaging was performed using a Nikon A1R HD25 and a 20x objective. Laser power and gain were optimized for each experiment, with the same setting applied to all treatments and controls.

The following antibodies were purchased from Cell Signaling Technologies: rabbit monoclonal antibodies against H3K27me3 (CST#9733), H3K27ac (CST#8173), H3K36me3 (CST#4909), H3K9me2 (CST#4658), H3K9ac (CST#9649), H3K4me1 (CST#9751), H3K4me3 (CST#5326), and 5mC (CST#28692), and mouse monoclonal antibodies against histone H3 (CST#14269) and 5hmC (CST#51660). A mouse monoclonal antibody against histone H3 was purchased from Proteintech (CAT#68345). The antibodies used in this study have been validated by the manufacturers to show no cross-reactivity with other core histones.

### zBrain pErk/tErk imaging

Larvae at 6 dpf are fixed overnight at 4°C in 4% paraformaldehyde (PFA) with 0.25% Triton X-100 in PBS (PBT), followed by three 5-minute washes in PBT. Larvae are then exposed to a bleaching solution containing 3% H_2_O_2_ and 1% KOH in water for 20-30min to remove pigmentation. For antigen retrieval, samples are incubated in 150 mM Tris-HCl (pH 9.0) at room temperature for 5 minutes, transferred to a 70°C water bath for 15 minutes, and washed twice in PBT. Permeabilization is performed using 0.05% Trypsin-EDTA on ice (typically 45 minutes for 6 dpf larvae), followed by two quick washes and a 10-minute wash in PBT. Blocking is done for 1 hour in PBT containing 2% normal goat serum (NGS), 1% BSA, and 1% DMSO. Primary antibody is diluted 1:500 in 1% BSA and 1% DMSO in PBT and incubated overnight at 4°C, followed by three 15-minute washes in PBT. Secondary antibody is applied under the same buffer conditions and incubated overnight at 4°C, with similar washes afterward. Samples are then mounted in low-melting-point agarose as close to the imaging surface as possible to reduce light scattering.

### Brain registration and analysis

The EpiBrain reference brains were generated by co-staining 6 dpf zebrafish with total Erk and either the anti-H3Tail antibody (Proteintech, #68345), antiH3Core antibody (CST, #14269), or PI. The resulting brains were registered to the zBrain tErk reference brain using the Computational Morphometry Toolkit (CMTK)^76–78^ with the command -awr 010203 -T 8 -X 52 -C 8 -G 80 -R 3 -A ’--accuracy 0.4’ -W ’--accuracy 1.6’. The *elavl3* brain image from the zBrain image database^33^ was used as the HCR reference brain. All reference brains were aligned with the zBrain Atlas^33^ to enable subsequent brain region analysis.

EpiBrain, zBrain, and HCR images are registered to their corresponding reference brains and analyzed using FIJI^79^ and Matlab (MathWorks, USA) by following an established method^33^. The Mann-Whitney U statistic Z score is calculated for each voxel to compare the treatment group with the control group, with significance threshold set at 0.05. The numbers of larvae images in each experimental group are documented in Supplementary Table 2. All image processing and computational analyses were conducted on a PC equipped with an AMD Ryzen Threadripper PRO 5955WX (16-Core) processor (Up to 4.0 GHz), a 128GB DDR4 memory, and a NVIDIA RTX A2000 6GB graphics card (Digital Storm).

Following zBrain ROI analysis, datasheets containing positive and negative signal for ROIs are generated. These regional signals were sorted in excel into forebrain (telencephalon, diencephalon, and ganglia), midbrain (mesencephalon), and hindbrain (rhombencephalon and spinal cord). The sum of the signal in each ROI was calculated for positive and negative signal.

### Pentylenetetrazol (PTZ) treatment

Pentylenetetrazol (PTZ; Cayman Chemicals, CAS# 54-95-5, Item No. 18682) was dissolved in DMSO at a stock concentration of 2 M. To induce brain-wide activity, n=20 zebrafish larvae were transferred into a 12-well plate inside a mesh-bottom insert at 5 dpf. Each well was pre-filled with 2 mL HEPES-buffered E3 medium. Treatment solutions were prepared by adding 15 µL of the 2 M PTZ stock to achieve a final concentration of 15 mM, or 15 µL DMSO as control. Following a 1-hour exposure, larvae were rinsed in fresh embryo media for 5 minutes before being fixed or maintained in embryo media overnight.

### Other chemical exposures

Fertilized zebrafish embryos were collected and transferred into 60 mm diameter, 20 mm deep Petri dishes at a density of 30 embryos per dish. Each dish contained 10 mL of HEPES-buffered E3 embryo medium. Chemical compounds were prepared as concentrated stock solutions in their respective solvents to ensure stability and stored at –20°C until use. Each compound was added to duplicate dishes at the desired final concentrations, while negative control dishes received equivalent volumes of the corresponding solvent. Dead embryos were removed at 1 and 2 dpf to maintain media quality and reduce contamination. For compound exposed only from 0 to 3 dpf, all surviving larvae were rinsed thoroughly at 3 dpf with fresh E3 medium and transferred to clean Petri dishes containing new E3 medium. Larvae were maintained under standard conditions until 6 dpf, at which point they were fixed in 4% paraformaldehyde (PFA) in PBS-T at 4°C.

UNC1999 (Cayman Chemical, CAS# 1431612-23-5) was applied at a final concentration of 20 µM from 0 to 3 dpf. GSK-J4 (Cayman Chemical, CAS# 1797983-09-5) was applied at a final concentration of 40 µM from 0 to 6 dpf. Valproic acid (VPA) (Cayman Chemical, CAS# 99-66-1) was applied at 10 µM from 0 to 6 dpf. Flumequine (Cayman Chemical, CAS# 42835-25-6) was applied at 25 µM from 0 to 3 dpf. Decitabine (MedChemExpress, CAS# 2353-33-5) was applied at 10 µM from 0 to 6 dpf. Bisphenol A (BPA) (Sigma-Aldrich, CAS# 80-05-7) was applied at 20 µM from 0 to 3 dpf. UNC0642 (MedChemExpress, CAS# 1481677-78-4) was applied at 10 µM from 0 to 6 dpf. C646 (MedChemExpress, CAS# 328968-36-1) was applied at 40 µM for 5 hours at 5 dpf. For ethanol treatment, 24 hpf embryos were exposed to 1% ethanol in E3 embryo medium for 2 hours and rinsed thoroughly in fresh embryo medium. Bobcat 339 (Cayman Chemicals, CAS# 2280037-51-4) was applied at 1 µM for 1 hour.

### CRISPR-Cas9 gene knockout

We designed three sets of sgRNAs targeting zebrafish genes to maximize the efficiency of F0 knockout, following established protocols^80–82^. sgRNA sequences were designed using CHOPCHOP^83^, and the full list of sgRNAs used in this study is provided in Supplemental Table 1. To increase the likelihood of generating loss-of-function mutations, sgRNAs were selected to target early coding exons, thereby promoting the introduction of premature stop codons. Each sgRNA template included the SP6 promoter sequence upstream of the target RNA sequence, followed by an overlap adapter complementary to the 5′ end of an 80 bp constant oligo, as described in an established protocol^84^.

Oligonucleotides were synthesized by Eurofins Scientific. Double-stranded DNA templates were generated by PCR using Phusion Hot-Start Flex DNA Polymerase (New England Biolabs) with gene-specific and constant oligos, then purified using the Zymo DNA Clean & Concentrator-5 Kit (Zymo Research). *In vitro* transcription of sgRNAs was performed using the MEGAscript SP6 Transcription Kit (Thermo Fisher Scientific), and the resulting sgRNAs were purified with the Zymo RNA Clean & Concentrator-5 Kit (Zymo Research). For zebrafish embryonic microinjection, adult male and female zebrafish were placed in mating cages overnight with a divider. The next morning, 1-cell-stage embryos were collected immediately before injection, after removing the divider to initiate spawning. The injection mixture was prepared by combining sgRNA with 2 µM Spy Cas9-NLS protein and 1X NEBuffer (New England Biolabs), and embryos were injected at the 1-cell stage.

### Social Isolation

Zebrafish were socially isolated starting at 24 hpf by placing individual embryos into alternating wells of a black 96-well plate. The plate was covered with a lid, and embryo viability and media volume were monitored daily to ensure proper development. Control fish were raised in pairs by placing two embryos per well into alternating wells of a black 96-well plate. At 6 dpf, larvae were removed from their wells, pooled, and immediately fixed for staining.

### RNA sequencing and data analysis

RNA samples were extracted from 6 dpf larvae treated with 15uM of PTZ or DMSO with an n of 20 and three replicates per condition, by following an established protocol^85^. Sample quality control was conducted using the Agilent 4150 Bioanalyzer, and only RNA samples with a RIN > 7 were selected for library preparation. For each sample, 200 ng of qualified RNA was used. mRNA was enriched from total RNA using oligo(dT)-attached magnetic beads, then fragmented with a fragmentation buffer. First-strand cDNA synthesis was performed using random N6 primers, followed by synthesis of double-stranded cDNA. The resulting DNA was end-repaired, 5’-phosphorylated, and modified to have a protruding ‘A’ at the 3’ end, allowing ligation of a bubble-shaped adapter with a complementary protruding ‘T’. Adapter-ligated fragments were amplified by PCR with specific primers, denatured to single strands, and converted into single-stranded circular DNA libraries using a bridged primer. Libraries were quality-checked and proceeded to sequencing only after passing QC. Libraries were further amplified using phi29 polymerase to generate DNA nanoballs (DNBs), each containing over 300 copies of the original DNA molecule. DNBs were loaded onto a patterned nanoarray, and 150 bp paired-end reads were generated using sequencing-by-synthesis technology. Sequencing was performed on the MGI T7 platform at a depth of 20M reads per sample.

Raw sequencing data quality was assessed using FastQC, which confirmed that the FASTQ files were of high quality. Reads were aligned to the NCBI RefSeq assembly of the GRCz11 transcriptome using the Salmon^86^ aligner. Aligned reads were summarized to the gene level using the Bioconductor package tximport^87^. To ensure reliable comparisons, we filtered out genes with low expression, retaining only those with an average log counts-per-million (logCPM) > 0. This reduced the dataset from 30,245 to 20,892 genes. Differential expression analysis was carried out using the edgeR package to fit a generalized linear model (GLM) to the count data and apply quasi-likelihood F-tests to make comparisons^88^. To account for unobserved sources of variability, particularly between control samples, we estimated a surrogate variable using the surrogate variable analysis (SVA) package^89^, and included it in the model. Differentially expressed genes were identified using a false discovery rate (FDR) threshold of < 0.05. All analyses were conducted in R version 4.3.1 under CentOS Linux 7 (Core).

### Larval and juvenile zebrafish brain dissection

Brains from 6 dpf and 25 dpf zebrafish were dissected for various downstream analyses. At the respective time points, fish were euthanized by immersion in ice-cold water for 15 minutes in Petri dishes. For dissection, each fish was positioned vertically with the dorsal side facing up in a pre-carved slot within dissection wax in a Petri dish. The lower half of the body was immobilized using Gorilla Glue to provide stability during the procedure. The Petri dish was filled with ice-cold phosphate-buffered saline (PBS) to prevent tissue desiccation. Brains were carefully extracted from the skull using the tip of a sharp syringe needle.

### Quantitative Real-Time PCR

To compare differential gene expressions between PTZ-treated and DMSO control fish, larvae were treated at 6 dpf and collected 1 hour post treatment, at which point their brains were dissected as described. Total RNA was extracted using the Quick-RNA Miniprep Kit (Zymo Research, Irvine, California) according to the manufacturer’s instructions. RNA concentrations were measured at 260 nm using a NanoDrop 1000 Spectrophotometer (Thermo Scientific, Waltham, Massachusetts). Extracted RNA was reverse-transcribed into cDNA using the SuperScript™ IV First-Strand Synthesis System (ThermoFisher Scientific, Waltham, Massachusetts) following the manufacturer’s protocol. Quantitative PCR (qPCR) was performed using SYBR Green qPCR Master Mix (GlpBio Technology Inc., Montclair, California) on a Bio-Rad CFX Connect Real-Time System (Bio-Rad, Hercules, California). Gene expression data were normalized to the housekeeping gene β-actin. All primers were designed using the NCBI Primer Design tool and synthesized by Eurofins Genomics (Louisville, Kentucky). Primer sequences are provided in Supplementary Table 3.

### Hybridization Chain Reaction (HCR)

Larvae were treated with PTZ for 1 hour at 5 dpf and collected at 6 dpf 24 hours after treatment. HCR was carried out using the HCR RNA-FISH v3.0 kit (Molecular Instruments). Probes targeting *elavl3*, *gadd45ba*, and *gadd45bb* were custom-designed and synthesized by Molecular Instruments (Supplementary Table 4). Brains stained by HCR were registered to the *elavl3* reference brain by co-staining using the *elavl3* probe.

### Histone and DNA Dot Blot Analysis

Brains from 25 dpf zebrafish were dissected for histone protein extraction. N=8 brains were collected for each experimental condition. Fish were treated with PTZ or DMSO at 24 dpf for one hour and allowed to recover for 24 hours prior to dissection. Each brain was sectioned into three regions: forebrain, midbrain, and hindbrain. The cutting locations for the different regions are shown in Supplementary Figure 11. The dissected regions were placed into separate tubes, and proteins were extracted via homogenization in RIPA buffer (Thermo Fisher, CAT# PI89900). The resulting lysates underwent acid extraction for histone proteins^90^. Histone concentrations were quantified using the Thermo Pierce BCA Kit (Thermo Fisher, CAT# A55865) with a microplate reader, and protein concentrations were normalized to 250 ng/µL using nuclease-free water. Dot blot analysis was performed on a 0.2 µm nitrocellulose membrane. Signals for histone modifications (H3K27ac and H3K9me2) were normalized to total H3 and compared between PTZ- and DMSO-treated groups.

Brains from 6 dpf zebrafish were dissected for genomic DNA extraction. N=20 brains were collected for each experimental condition. Fish were treated with PTZ or DMSO at 5 dpf and allowed to recover for 24 hours before dissection. Brains were bisected into two regions: midbrain and hindbrain. The forebrain was excluded due to difficulties in consistently obtaining larval forebrain tissue. These sections were placed into individual tubes, and genomic DNA was extracted using the Zymo Quick-DNA MiniPrep Kit (Zymo Research, CAT# D3024). DNA concentrations were measured at 260 nm using a NanoDrop 1000 Spectrophotometer (Thermo Scientific, Waltham, Massachusetts), and adjusted to 25 ng/µL with nuclease-free water. Samples were normalized to total DNA concentration. Dot blot analysis was conducted on a nylon membrane following an established protocol^91^.

### Locomotor assay

Locomotor activity was assessed by video tracking of live 6 dpf larvae placed individually (1 per well) in glass 9-well depression plates, each containing approximately 3 mL of fresh, 26°C conditioned water. The plates were housed in a surround-lit video enclosure isolated from external stimuli. Top-down video recordings were captured at 30 frames per second (fps) for 11 minutes immediately following larval distribution. Horizontal positional data were extracted from the videos using Bonsai^92^, a visual programming suite, at a resolution of one position per frame. The first minute of each recording was excluded using Python to control for variability during acclimation. Python scripts were also used to compile locomotor metrics for each individual larva and treatment group. The resulting locomotor data consisted of total distance traveled per larva and frame-by-frame movement data. Using a 1/30 second interval, per-frame displacement was used to approximate the average horizontal velocity for each larva.

### Statistical analysis

Graphs were generated using GraphPad Prism. Two-tailed Student’s *t* test was used to analyze data when there were two groups. For analysis of multiple groups, normal distribution of datasets was examined using Shapiro-Wilk test, and if more than half the data was normally distributed and standard deviations fell within the variance ratio, one-way analysis of variance (ANOVA) assuming gaussian distribution and equal standard deviation was performed, using Tukey’s or Šídák’s multiple comparison test for post hoc analysis. *P* values less than 0.05 were considered significant.

## ACKNOWLEDGEMENTS

We thank the University of Washington Office of Comparative Medicine for providing zebrafish husbandry support. This work was supported by the National Institute of Environmental Health Sciences (NIEHS) of the NIH under the award number R00ES031050 and the National Institute on Drug Abuse (NIDA) of the NIH under the award number P30DA048736. The content is solely the responsibility of the authors and does not necessarily represent the official views of the NIH.

## AUTHOR CONTRIBUTIONS

M.J. designed and conducted the experiments and analyzed data. A.E. and K.S.G. assisted in conducting experiments. J.W.M and T.K.B analyzed RNA-seq data. Y.G. conceived the study and interpreted the data. M.J. and Y.G. wrote the manuscript. All authors contributed meaningful insights during discussions and reviewed and approved the final version of the manuscript.

## DATA AND MATERIALS AVAILABILITY

All data needed to evaluate the conclusions in the paper are present in the paper and/or the Supplementary Materials.

## REFERENCES

1 Oldroyd, B. P. & Yagound, B. The role of epigenetics, particularly DNA methylation, in the evolution of caste in insect societies. Philos Trans R Soc Lond B Biol Sci 376, 20200115 (2021). PMC8059649.

2 Opachaloemphan, C., Yan, H., Leibholz, A., Desplan, C. & Reinberg, D. Recent Advances in Behavioral (Epi)Genetics in Eusocial Insects. Annu Rev Genet 52, 489–510 (2018). PMC6445553.

3 Sieber, K. R., Dorman, T., Newell, N. & Yan, H. (Epi)Genetic Mechanisms Underlying the Evolutionary Success of Eusocial Insects. Insects 12 (2021). PMC8229086.

4 Weiner, S. A. & Toth, A. L. Epigenetics in social insects: a new direction for understanding the evolution of castes. Genet Res Int 2012, 609810 (2012). PMC3335566.

5 Fagiolini, M., Jensen, C. L. & Champagne, F. A. Epigenetic influences on brain development and plasticity. Curr Opin Neurobiol 19, 207–212 (2009). PMC2745597.

6 Feng, J., Fouse, S. & Fan, G. Epigenetic regulation of neural gene expression and neuronal function. Pediatr Res 61, 58R–63R (2007).

7 Nelson, E. D. & Monteggia, L. M. Epigenetics in the mature mammalian brain: effects on behavior and synaptic transmission. Neurobiol Learn Mem 96, 53–60 (2011). PMC3463371.

8 Pizzimenti, C. L. & Lattal, K. M. Epigenetics and memory: causes, consequences and treatments for post-traumatic stress disorder and addiction. Genes Brain Behav 14, 73–84 (2015). PMC4526190.

9. Diaz, L., et al. Butyrate rescues chlorpyrifos-induced social deficits through inhibition of class I histone deacetylases. bioRxiv (2025).

10 Geng, Y. et al. Top2a promotes the development of social behavior via PRC2 and H3K27me3. Sci Adv 8, eabm7069 (2022). PMC9683714.

11 Rudenko, A. & Tsai, L. H. Epigenetic modifications in the nervous system and their impact upon cognitive impairments. Neuropharmacology 80, 70–82 (2014).

12 Nayak, M. et al. Epigenetic signature in neural plasticity: the journey so far and journey ahead. Heliyon 8, e12292 (2022). PMC9798197.

13 Kuehner, J. N., Bruggeman, E. C., Wen, Z. & Yao, B. Epigenetic Regulations in Neuropsychiatric Disorders. Front Genet 10, 268 (2019). PMC6458251.

14 Tsankova, N., Renthal, W., Kumar, A. & Nestler, E. J. Epigenetic regulation in psychiatric disorders. Nat Rev Neurosci 8, 355–367 (2007).

15 Abdolmaleky, H. M., Zhou, J. R. & Thiagalingam, S. Cataloging recent advances in epigenetic alterations in major mental disorders and autism. Epigenomics 13, 1231–1245 (2021). PMC8738978.

16 Moos, W. H. et al. Epigenetic Treatment of Neuropsychiatric Disorders: Autism and Schizophrenia. Drug Dev Res 77, 53–72 (2016).

17 Richetto, J. & Meyer, U. Epigenetic Modifications in Schizophrenia and Related Disorders: Molecular Scars of Environmental Exposures and Source of Phenotypic Variability. Biol Psychiatry 89, 215–226 (2021).

18 Tremblay, M. W. & Jiang, Y. H. DNA Methylation and Susceptibility to Autism Spectrum Disorder. Annu Rev Med 70, 151–166 (2019). PMC6597259.

19 Nestler, E. J., Pena, C. J., Kundakovic, M., Mitchell, A. & Akbarian, S. Epigenetic Basis of Mental Illness. Neuroscientist 22, 447–463 (2016). PMC4826318.

20 Kaplan, G., Xu, H., Abreu, K. & Feng, J. DNA Epigenetics in Addiction Susceptibility. Front Genet 13, 806685 (2022). PMC8821887.

21 Nielsen, D. A., Utrankar, A., Reyes, J. A., Simons, D. D. & Kosten, T. R. Epigenetics of drug abuse: predisposition or response. Pharmacogenomics 13, 1149–1160 (2012). PMC3463407.

22 Walker, D. M. & Nestler, E. J. Neuroepigenetics and addiction. Handb Clin Neurol 148, 747–765 (2018). PMC5868351.

23 Sng, J. & Meaney, M. J. Environmental regulation of the neural epigenome. Epigenomics 1, 131–151 (2009).

24. Wang, F., Pan, F., Tang, Y. & Huang, J. H. Editorial: Early Life Stress-Induced Epigenetic Changes Involved in Mental Disorders. Front Genet 12, 684844 (2021). PMC8320347.

25 Rahman, M. F. & McGowan, P. O. Cell-type-specific epigenetic effects of early life stress on the brain. Transl Psychiatry 12, 326 (2022). PMC9365848.

26 Cheng, Z., Su, J., Zhang, K., Jiang, H. & Li, B. Epigenetic Mechanism of Early Life Stress-Induced Depression: Focus on the Neurotransmitter Systems. Front Cell Dev Biol 10, 929732 (2022). PMC9294154.

27 Unternaehrer, E. et al. Dynamic changes in DNA methylation of stress-associated genes (OXTR, BDNF ) after acute psychosocial stress. Transl Psychiatry 2, e150 (2012). PMC3432191.

28 Fu, J. M. et al. Rare coding variation provides insight into the genetic architecture and phenotypic context of autism. Nat Genet 54, 1320–1331 (2022). PMC9653013.

29 Satterstrom, F. K. et al. Large-Scale Exome Sequencing Study Implicates Both Developmental and Functional Changes in the Neurobiology of Autism. Cell 180, 568–584 e523 (2020). PMC7250485.

30 Crews, D., Gillette, R., Miller-Crews, I., Gore, A. C. & Skinner, M. K. Nature, nurture and epigenetics. Mol Cell Endocrinol 398, 42–52 (2014). PMC4300943.

31 Moore, D. S. Behavioral epigenetics. Wiley Interdiscip Rev Syst Biol Med 9 (2017).

32 Tammen, S. A., Friso, S. & Choi, S. W. Epigenetics: the link between nature and nurture. Mol Aspects Med 34, 753–764 (2013). PMC3515707.

33 Randlett, O. et al. Whole-brain activity mapping onto a zebrafish brain atlas. Nat Methods 12, 1039–1046 (2015). PMC4710481.

34 Mai, H. et al. Whole-body cellular mapping in mouse using standard IgG antibodies. Nat Biotechnol 42, 617–627 (2024). PMC11021200.

35 Kruidenier, L. et al. A selective jumonji H3K27 demethylase inhibitor modulates the proinflammatory macrophage response. Nature 488, 404–408 (2012). PMC4691848.

36 Moshi, J. M., Ummelen, M., Broers, J. L. V., Ramaekers, F. C. S. & Hopman, A. H. N. Impact of antigen retrieval protocols on the immunohistochemical detection of epigenetic DNA modifications. Histochem Cell Biol 159, 513–526 (2023). PMC10247850.

37 Tang, X., Falls, D. L., Li, X., Lane, T. & Luskin, M. B. Antigen-retrieval procedure for bromodeoxyuridine immunolabeling with concurrent labeling of nuclear DNA and antigens damaged by HCl pretreatment. J Neurosci 27, 5837–5844 (2007). PMC6672250.

38 Wang, K. et al. TSA-PACT: a method for tissue clearing and immunofluorescence staining on zebrafish brain with improved sensitivity, specificity and stability. Cell Biosci 13, 97 (2023). PMC10223841.

39 Lee, E. et al. ACT-PRESTO: Rapid and consistent tissue clearing and labeling method for 3-dimensional (3D) imaging. Sci Rep 6, 18631 (2016). PMC4707495.

40 Sasaki, K. et al. Effects of denaturation with HCl on the immunological staining of bromodeoxyuridine incorporated into DNA. Cytometry 9, 93–96 (1988).

41 Liang, H., Wu, X., Yalowich, J. C. & Hasinoff, B. B. A three-dimensional quantitative structure-activity analysis of a new class of bisphenol topoisomerase IIalpha inhibitors. Mol Pharmacol 73, 686–696 (2008).

42 Di Ciaula, A. & Portincasa, P. Diet and Contaminants: Driving the Rise to Obesity Epidemics? Curr Med Chem 26, 3471–3482 (2019).

43 Janesick, A. & Blumberg, B. Obesogens, stem cells and the developmental programming of obesity. Int J Androl 35, 437–448 (2012). PMC3358413.

44 Desai, M., Ferrini, M. G., Han, G., Jellyman, J. K. & Ross, M. G. In vivo maternal and in vitro BPA exposure effects on hypothalamic neurogenesis and appetite regulators. Environ Res 164, 45–52 (2018). PMC8085909.

45 Richter, C. A. et al. In vivo effects of bisphenol A in laboratory rodent studies. Reprod Toxicol 24, 199–224 (2007). PMC2151845.

46 Rubin, B. S. & Soto, A. M. Bisphenol A: Perinatal exposure and body weight. Mol Cell Endocrinol 304, 55–62 (2009). PMC2817931.

47. Karin, S. & Bondesson, M. Bisphenol A and its Analogues Alter Appetite Control in Zebrafish. *bioRxiv* (2024).

48 Sugiyama, K. I., Kinoshita, M., Gruz, P., Kasamatsu, T. & Honma, M. Bisphenol-A reduces DNA methylation after metabolic activation. Genes Environ 44, 20 (2022). PMC9316663.

49 Cervera-Juanes, R., Wilhelm, L. J., Park, B., Grant, K. A. & Ferguson, B. Alcohol-dose-dependent DNA methylation and expression in the nucleus accumbens identifies coordinated regulation of synaptic genes. Transl Psychiatry 7, e994 (2017). PMC5545731.

50 Bonsch, D., Lenz, B., Reulbach, U., Kornhuber, J. & Bleich, S. Homocysteine associated genomic DNA hypermethylation in patients with chronic alcoholism. J Neural Transm (Vienna) 111, 1611–1616 (2004).

51 Portales-Casamar, E. et al. DNA methylation signature of human fetal alcohol spectrum disorder. Epigenetics Chromatin 9, 25 (2016). PMC4926300.

52 Pinheiro-da-Silva, J. & Luchiari, A. C. Embryonic ethanol exposure on zebrafish early development. Brain Behav 11, e02062 (2021). PMC8213935.

53 Komada, M. et al. Mechanisms underlying neuro-inflammation and neurodevelopmental toxicity in the mouse neocortex following prenatal exposure to ethanol. Sci Rep 7, 4934 (2017). PMC5504035.

54 Shetty, A. K., Burrows, R. C. & Phillips, D. E. Alterations in neuronal development in the substantia nigra pars compacta following in utero ethanol exposure: immunohistochemical and Golgi studies. Neuroscience 52, 311–322 (1993).

55 Kuehner, J. N. et al. Social defeat stress induces genome-wide 5mC and 5hmC alterations in the mouse brain. G3 (Bethesda) 13 (2023). PMC10411578.

56 Nestler, E. J. & Luscher, C. The Molecular Basis of Drug Addiction: Linking Epigenetic to Synaptic and Circuit Mechanisms. Neuron 102, 48–59 (2019). PMC6587180.

57 Stewart, A. F., Fulton, S. L. & Maze, I. Epigenetics of Drug Addiction. Cold Spring Harb Perspect Med 11 (2021). PMC7967246.

58 Toth, M. Epigenetic Neuropharmacology: Drugs Affecting the Epigenome in the Brain. Annu Rev Pharmacol Toxicol 61, 181–201 (2021). PMC9915026.

59 Turrini, L. et al. Optical mapping of neuronal activity during seizures in zebrafish. Sci Rep 7, 3025 (2017). PMC5465210.

60 Carpenter, M. D. et al. Nr4a1 suppresses cocaine-induced behavior via epigenetic regulation of homeostatic target genes. Nat Commun 11, 504 (2020). PMC6981219.

61 Ma, D. K. et al. Neuronal activity-induced Gadd45b promotes epigenetic DNA demethylation and adult neurogenesis. Science 323, 1074–1077 (2009). PMC2726986.

62 Hrvatin, S. et al. Single-cell analysis of experience-dependent transcriptomic states in the mouse visual cortex. Nat Neurosci 21, 120–129 (2018). PMC5742025.

63 Moskalev, A. A. et al. Gadd45 proteins: relevance to aging, longevity and age-related pathologies. Ageing Res Rev 11, 51–66 (2012). PMC3765067.

64 Hanotel, J. et al. The Prdm13 histone methyltransferase encoding gene is a Ptf1a-Rbpj downstream target that suppresses glutamatergic and promotes GABAergic neuronal fate in the dorsal neural tube. Dev Biol 386, 340–357 (2014).

65 Pollina, E. A. et al. A NPAS4-NuA4 complex couples synaptic activity to DNA repair. Nature 614, 732–741 (2023). PMC9946837.

66 Spaapen, F. et al. The immediate early gene product EGR1 and polycomb group proteins interact in epigenetic programming during chondrogenesis. PLoS One 8, e58083 (2013). PMC3590300.

67 Sun, Z. et al. EGR1 recruits TET1 to shape the brain methylome during development and upon neuronal activity. Nat Commun 10, 3892 (2019). PMC6715719.

68. Yin, L., et al. Elevated EGR1 Binding at Enhancers in Excitatory Neurons Correlates with Neuronal Subtype-Specific Epigenetic Regulation. bioRxiv (2024). PMC11601525.

69 Mendes, K. et al. The epigenetic pioneer EGR2 initiates DNA demethylation in differentiating monocytes at both stable and transient binding sites. Nat Commun 12, 1556 (2021). PMC7946903.

70 Tsai, M. H., Wuputra, K., Lin, Y. C., Lin, C. S. & Yokoyama, K. K. Multiple functions of the histone chaperone Jun dimerization protein 2. Gene 590, 193–200 (2016). PMC6639046.

71 Dirks, R. M. & Pierce, N. A. Triggered amplification by hybridization chain reaction. Proc Natl Acad Sci U S A 101, 15275–15278 (2004). PMC524468.

72 Chen, A. et al. QRFP and Its Receptors Regulate Locomotor Activity and Sleep in Zebrafish. J Neurosci 36, 1823–1840 (2016). PMC4748070.

73 Barker, A. J. & Baier, H. Sensorimotor decision making in the zebrafish tectum. Curr Biol 25, 2804–2814 (2015).

74 Portugues, R., Feierstein, C. E., Engert, F. & Orger, M. B. Whole-brain activity maps reveal stereotyped, distributed networks for visuomotor behavior. Neuron 81, 1328–1343 (2014). PMC4448760.

75 Geng, Y. & Peterson, R. T. Rapid Mounting of Zebrafish Larvae for Brain Imaging. Zebrafish 18, 376–379 (2021). PMC8716516.

76 Ostrovsky, A., Cachero, S. & Jefferis, G. Clonal analysis of olfaction in Drosophila: image registration. Cold Spring Harb Protoc 2013, 347–349 (2013).

77 Jefferis, G. S. et al. Comprehensive maps of Drosophila higher olfactory centers: spatially segregated fruit and pheromone representation. Cell 128, 1187–1203 (2007). PMC1885945.

78 Rohlfing, T. & Maurer, C. R., Jr. Nonrigid image registration in shared-memory multiprocessor environments with application to brains, breasts, and bees. IEEE Trans Inf Technol Biomed 7, 16–25 (2003).

79. Schindelin, J., et al. Fiji: an open-source platform for biological-image analysis. Nat Methods 9, 676–682 (2012). PMC3855844.

80 Kim, B. H. & Zhang, G. Generating Stable Knockout Zebrafish Lines by Deleting Large Chromosomal Fragments Using Multiple gRNAs. G3 (Bethesda) 10, 1029–1037 (2020). PMC7056962.

81 Kroll, F. et al. A simple and effective F0 knockout method for rapid screening of behaviour and other complex phenotypes. Elife 10 (2021). PMC7793621.

82 Wu, R. S. et al. A Rapid Method for Directed Gene Knockout for Screening in G0 Zebrafish. Dev Cell 46, 112–125 e114 (2018).

83 Montague, T. G., Cruz, J. M., Gagnon, J. A., Church, G. M. & Valen, E. CHOPCHOP: a CRISPR/Cas9 and TALEN web tool for genome editing. Nucleic Acids Res 42, W401–407 (2014). PMC4086086.

84 Gagnon, J. A. et al. Efficient mutagenesis by Cas9 protein-mediated oligonucleotide insertion and large-scale assessment of single-guide RNAs. PLoS One 9, e98186 (2014). PMC4038517.

85 Peterson, S. M. & Freeman, J. L. RNA isolation from embryonic zebrafish and cDNA synthesis for gene expression analysis. J Vis Exp (2009). PMC3152201.

86 Patro, R., Duggal, G., Love, M. I., Irizarry, R. A. & Kingsford, C. Salmon provides fast and bias-aware quantification of transcript expression. Nat Methods 14, 417–419 (2017). PMC5600148.

87 Soneson, C., Love, M. I. & Robinson, M. D. Differential analyses for RNA-seq: transcript-level estimates improve gene-level inferences. F1000Res 4, 1521 (2015). PMC4712774.

88 Lund, S. P., Nettleton, D., McCarthy, D. J. & Smyth, G. K. Detecting differential expression in RNA-sequence data using quasi-likelihood with shrunken dispersion estimates. Stat Appl Genet Mol Biol 11 (2012).

89 Leek, J. T. & Storey, J. D. Capturing heterogeneity in gene expression studies by surrogate variable analysis. PLoS Genet 3, 1724–1735 (2007). PMC1994707.

90. Holt, M. V., Wang, T. & Young, N. L. Expeditious Extraction of Histones from Limited Cells or Tissue Samples and Quantitative Top-Down Proteomic Analysis. Curr Protoc 1, e26 (2021). PMC8991418.

91 Jia, Z. et al. A 5-mC Dot Blot Assay Quantifying the DNA Methylation Level of Chondrocyte Dedifferentiation In Vitro. J Vis Exp (2017). PMC5607994.

92 Lopes, G. et al. Bonsai: an event-based framework for processing and controlling data streams. Front Neuroinform 9, 7 (2015). PMC4389726.

