## Supplementary Materials for "EpiBrain: the brain’s epigenetic landscape in a snapshot"

**This PDF file includes:**

- Supplementary Figures 1 to 11
- Supplementary Tables 1 to 4 Legends
- Supplementary Data 1 to 5 Legends

### SUPPLEMENTARY FIGURES

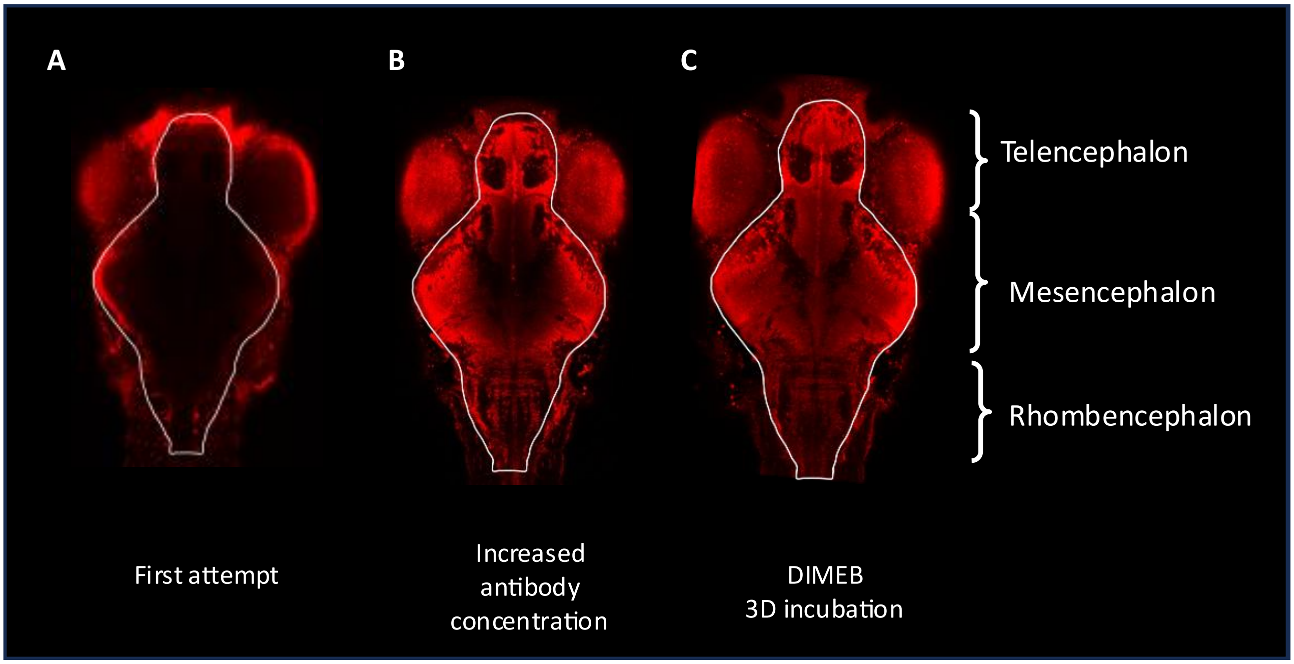


**Supplementary Figure 1.**

(**A**) The first attempt at histone H3 staining of 6 dpf larval zebrafish brain by following a base protocol was unsuccessful. (**B**) Increasing the amount of anti-H3 antibody from a 1:500 dilution to a 3:500 dilution significantly improved staining signal, but still left the center of the brain insufficiently stained. (**C**) The addition of heptakis(2,6-di-O-methyl)-β-cyclodextrin (DIMEB) and increased antibody incubation time (3 days) significantly improved antibody penetration and resulted in homogeneous staining of the larval brain.

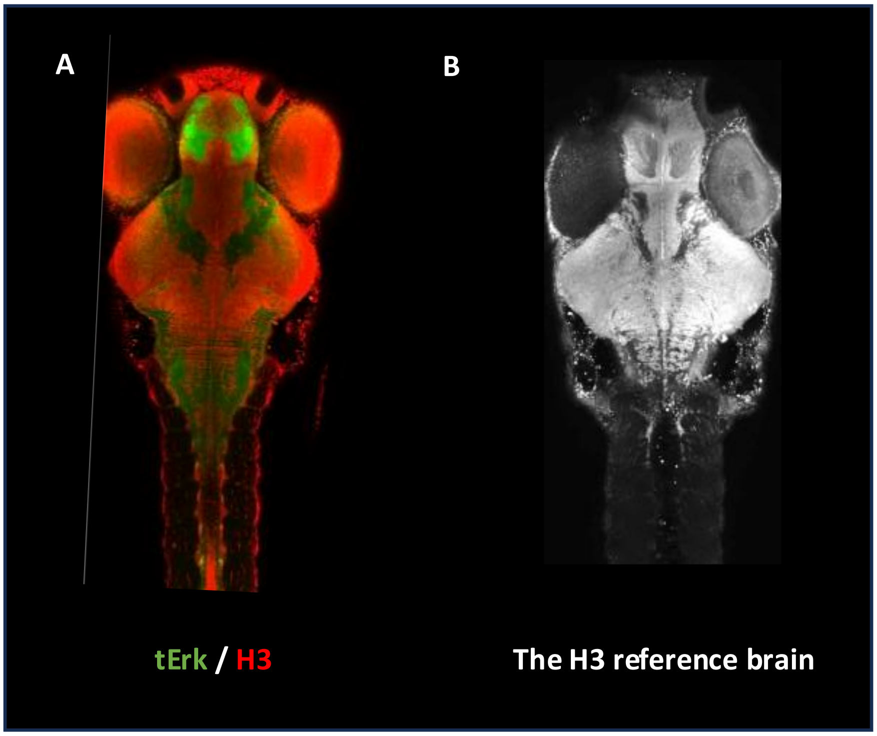


**Supplementary Figure 2.**

(**A**) A wild-type larvae brain counterstained for histone H3 and total Erk (tErk), and registered to the zBrain altas' tErk template brain. (**B**) A cross-sectional image of the histone H3 template brain for EpiBrain image registration.

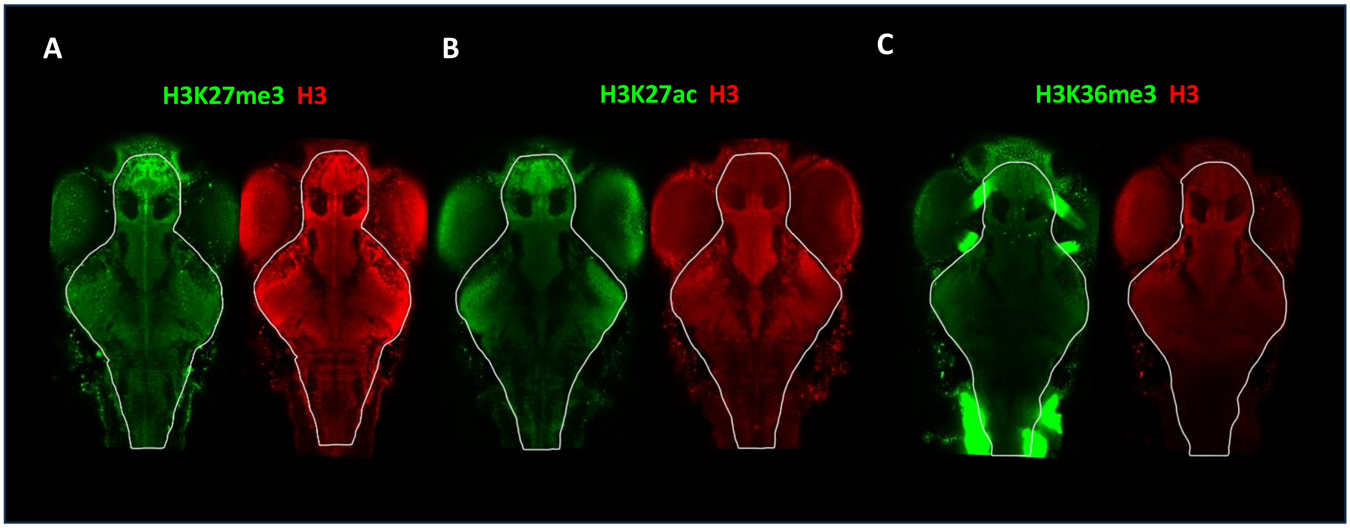


**Supplementary Figure 3.**

(**A-C**) Cross-sectional images of histone H3 counterstained with H3K27me3 (A), H3K27ac (B), and H3K36me3 (C).

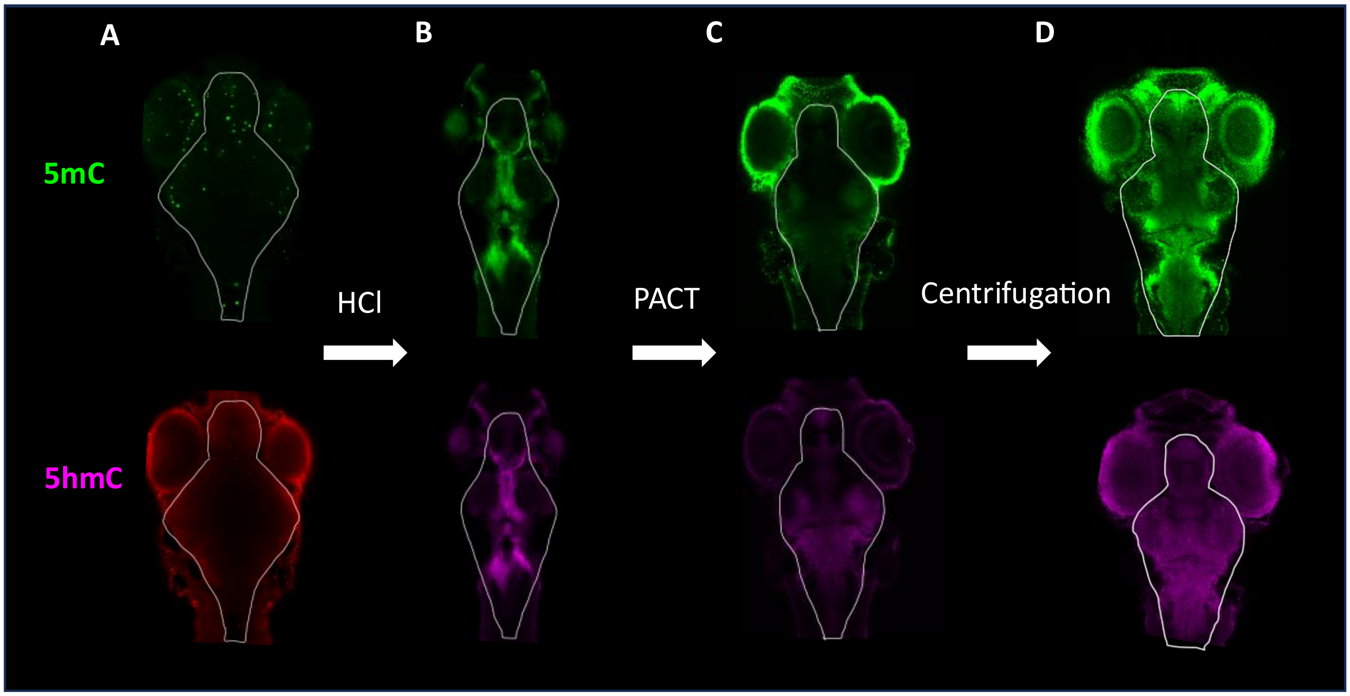


**Supplementary Figure 4.**

(**A**) The EpiBrain-H3(K27/K36) protocol was unsuccessful at staining the DNA modifications 5mC (top row) and 5hmC (bottom row). (**B**) HCl treatment effectively boosted staining signal, but caused significant changes in brain morphology. (**C**) Employing passive CLARITY technique (PACT) prior to HCl treatment helped maintain normal tissue morphology. However, antibody penetration remained suboptimal following these procedures. (**D**) Centrifugal pressure significantly boosted antibody penetration and helped achieve homogeneous staining.

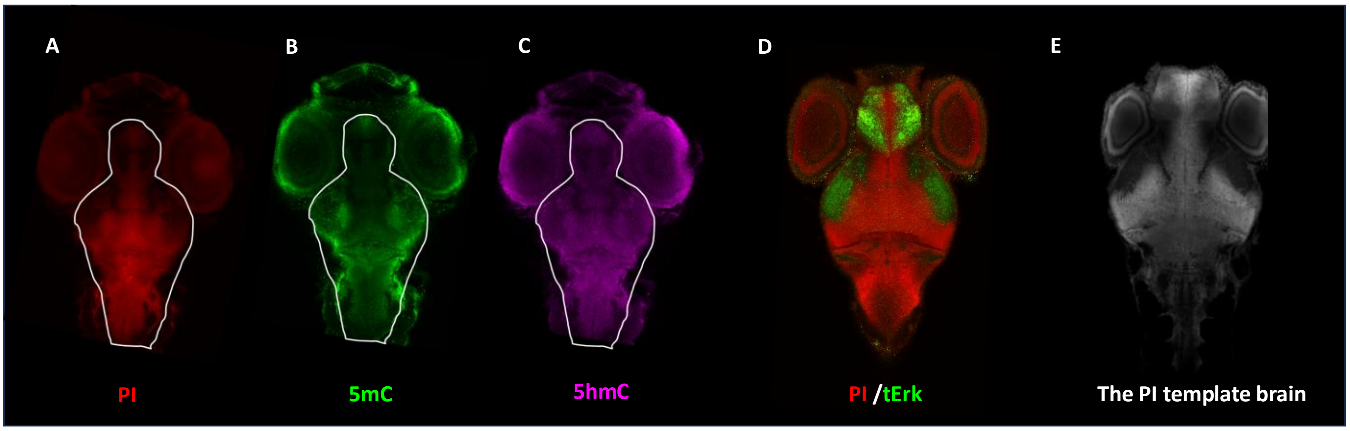


**Supplementary Figure 5.**

(**A-C**) Propidium iodide (PI) (A) showed excellent compatibility with 5mC (B) and 5hmC (C) co-staining. (**D**) A wild-type larvae brain counterstained for PI and tErk, and registered to the zBrain altas' tErk template brain. (**E**) A cross-sectional image of the PI template brain for EpiBrain image registration, created by registering a tErk/PI counterstained brain to the tErk template brain.

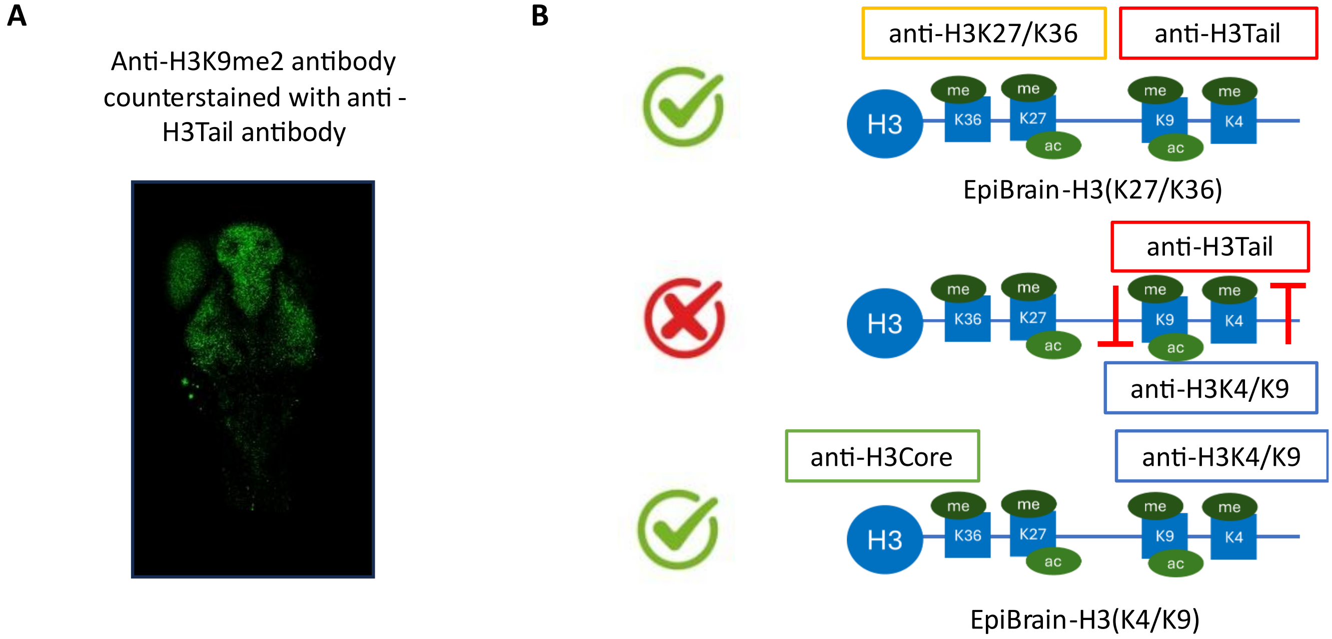


**Supplementary Figure 6.**

(**A**) The anti-H3K9me2 antibody did not successfully counterstain with the anti-H3Tail antibody. (**B**) A schematic illustration of the hypothesized reason for why the anti-H3Tail antibody failed to counterstain with the anti-H3K9me2 antibody. The anti-H3Tail antibody binds to the N-terminus (tail region) of histone H3, which does not overlap with the bindings of anti-H3K27 and anti-H3K36 antibodies, enabling it to be used for the EpiBrain-H3(K27/K36) protocol. However, anti-H3K4 and anti-H3K9 antibodies directly compete with the binding of anti-H3Tail antibody at the histone tail region, therefore requiring an anti-H3Core antibody which targets the core histone region (C-terminus) for counterstaining.

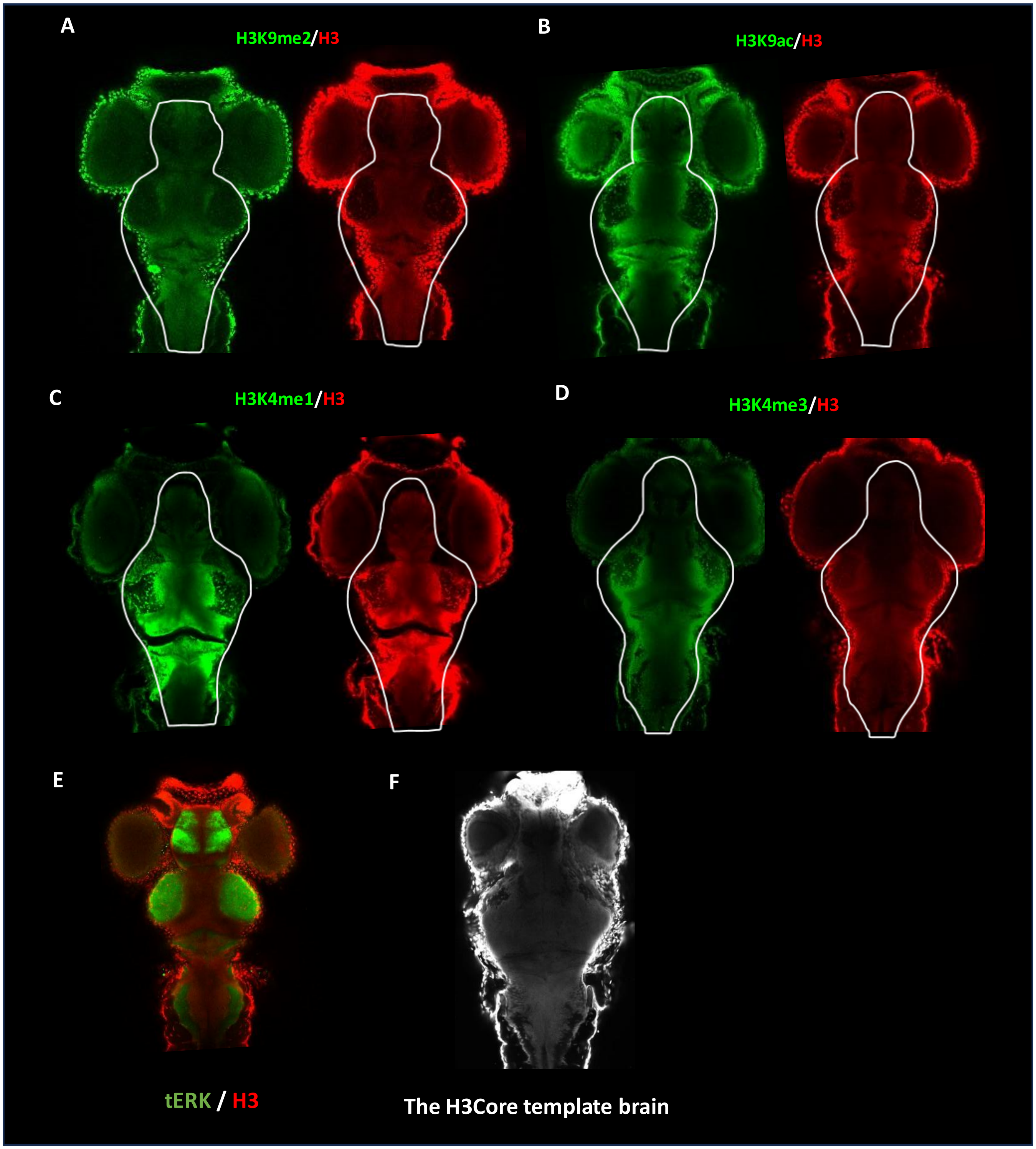


**Supplementary Figure 7.**

(**A-D**) The EpiBrain-H3(K4/K9) protocol successfully counterstained an anti-H3Core antibody with anti-H3K9me2 (A), anti-H3K9ac (B), anti-H3K4me1 (C), and anti-H3K4me3 (D) antibodies. (**E**) A wild-type larvae brain counterstained for histone H3 using the anti-H3Core antibody and total Erk (tErk), and registered to the zBrain altas' tErk template brain. (**B**) A cross-sectional image of the histone H3Core template brain for EpiBrain-H3(K4/K9) image registration.

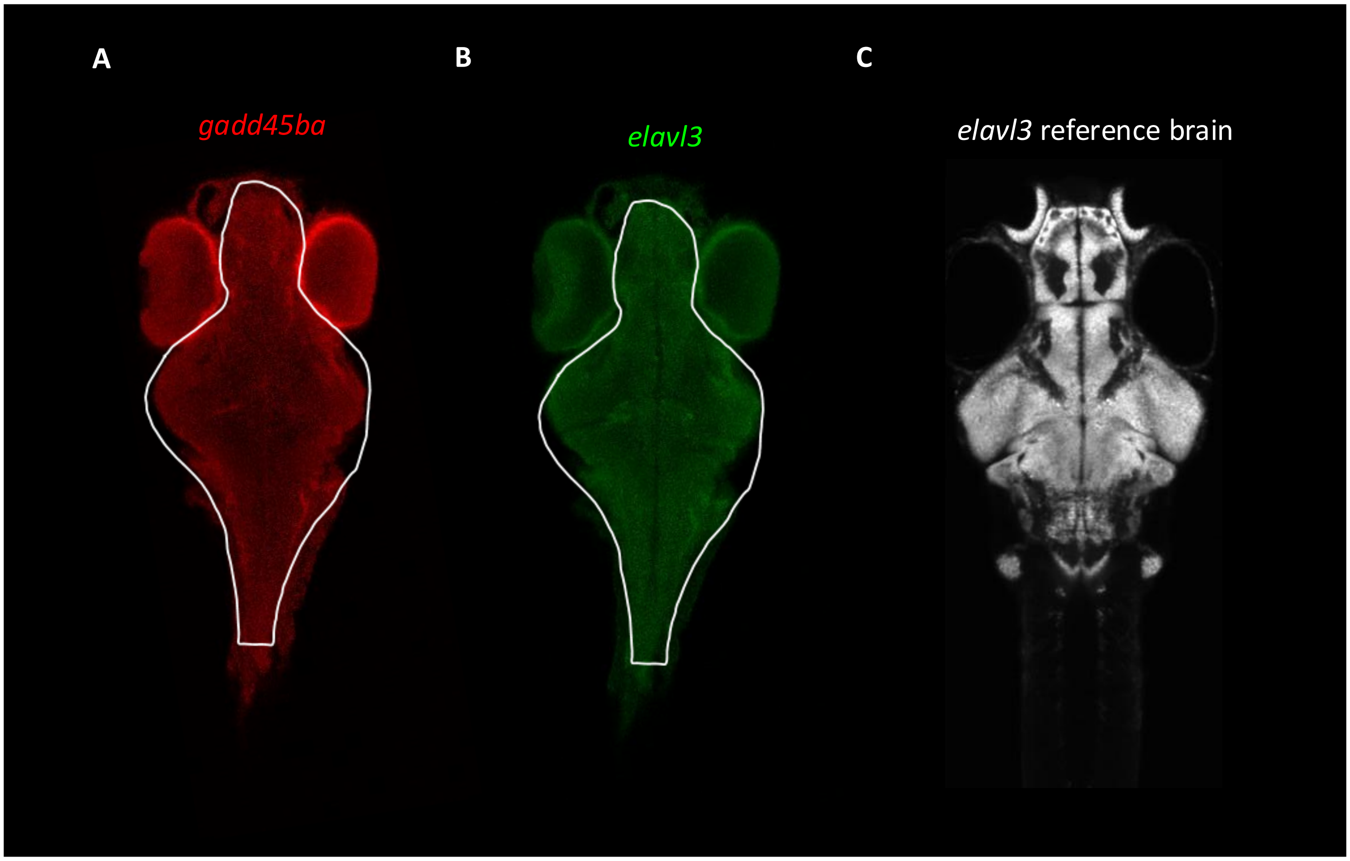


**Supplementary Figure 8.**

(**A**) HCR detected *gadd45ba* expression throughout the 6 dpf larval brain. (**B**) Co-staining of *elavl3* using HCR. (**C**) The *gadd45ba*/*elavl3* counterstained images were registered to an *elavl3* reference brain acquired from the zBrain atlas to compare gene expression with or without PTZ treatment in various regions of the brain.

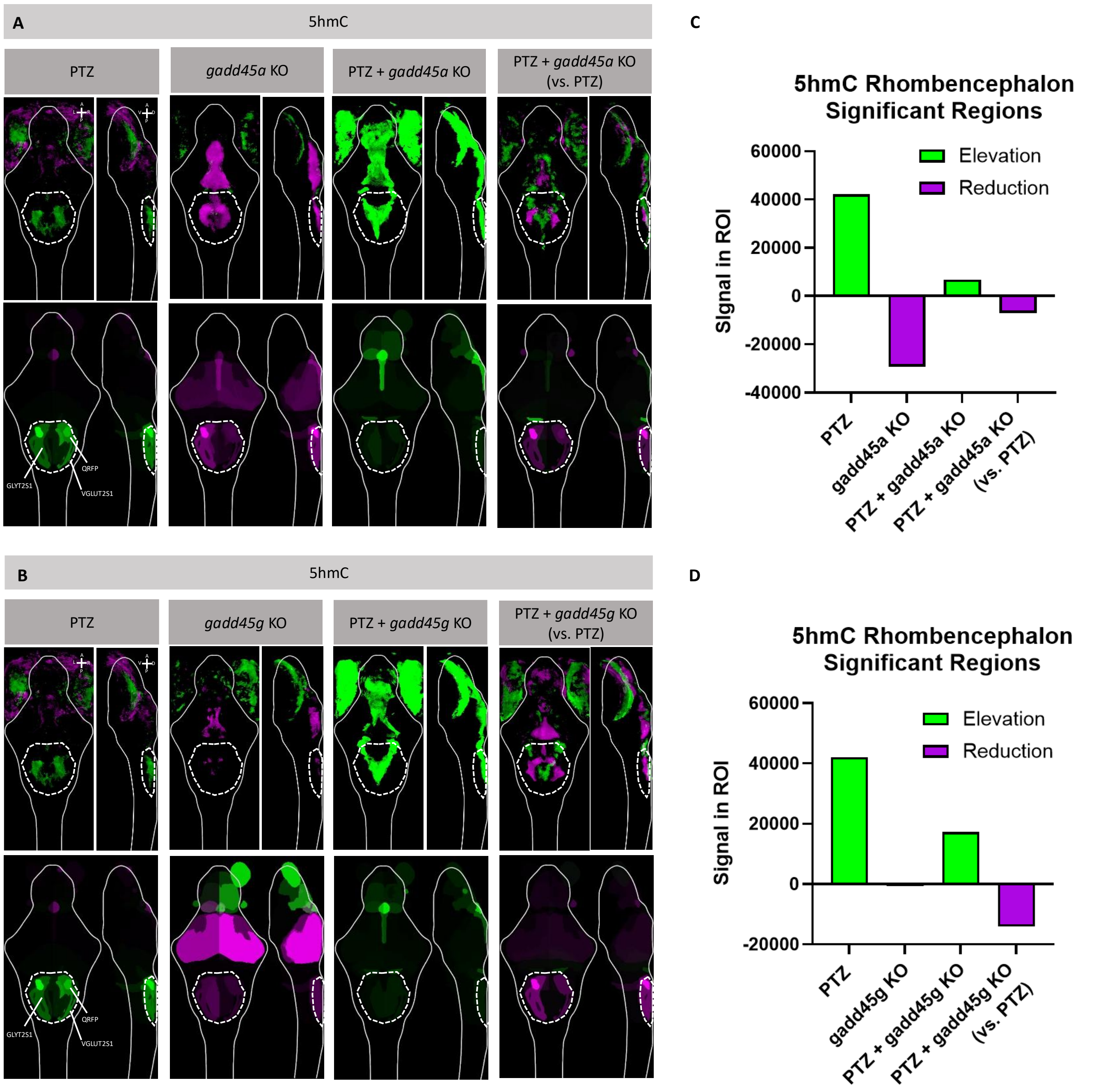


**Supplementary Figure 9.**

(**A-B**) EpiBrain-DNA imaging and analysis detected changes in 5hmC in various brain regions following PTZ treatment and *gadd45a* (A) or *gadd45g* (B) KOs. Green: elevation. Magenta: reduction. PTZ: 1-hr PTZ treatment at 5 dpf; analysis was conducted by comparing to DMSO control; re-adaptation of Figure 3L. *gadd45a/g* KO: F0 CRISPR/Cas9 double-KO of *gadd45aa* and *gadd45ab* (*gadd45a* KO) (A) or *gadd45ga* and *gadd45gb* (*gadd45g* KO) (B); analysis was conducted by comparing to wild-type (WT) control. PTZ + *gadd45a/g* KO: *gadd45a/g* KO larvae exposed to PTZ for 1-hr at 5 dpf; analysis was conducted by comparing to WT control. PTZ + *gadd45a/g* KO (vs. PTZ): *gadd45a/g* KO larvae exposed to PTZ for 1-hr at 5 dpf; analysis was conducted by comparing to WT control exposed to PTZ for 1-hr at 5 dpf. Brain regions are identified based on alignment with the zBrain atlas following brain registration. The top differentially regulated brain regions are labeled in the images. Raw analysis results for all differentially regulated brain regions and their signal intensities in images shown in Supplementary Figure 9 are summarized in Supplementary Data 5. GLYT2S1: glyt2 stripe 1. QRFP: qrfp neuron cluster sparse. VGLUT2S1: vglut2 stripe 1. (**C-D**) Bar plots showing summed 5hmC signal changes in rhombencephalon regions of interest (ROIs; white dashed circles in Supplementary Figures 9A & 9B) following PTZ exposure, *gadd45a/g* KO, combined PTZ + *gadd45a/g* KO, and PTZ + *gadd45a/g* KO relative to PTZ alone. Rhombencephalon regions were first identified based on significant 5hmC changes in each condition in the *gadd45b* KO experiment (Figure 4D), and only regions consistently detected across all comparisons were included in the analysis. For each condition, 5hmC signal was summed across these shared ROIs and plotted, with positive values indicating elevation and negative values indicating reduction in 5hmC signal. The analysis reveals PTZ-induced increases in 5hmC (PTZ) that are partially reversed by *gadd45a/g* knockout (PTZ + *gadd45a/g* KO).

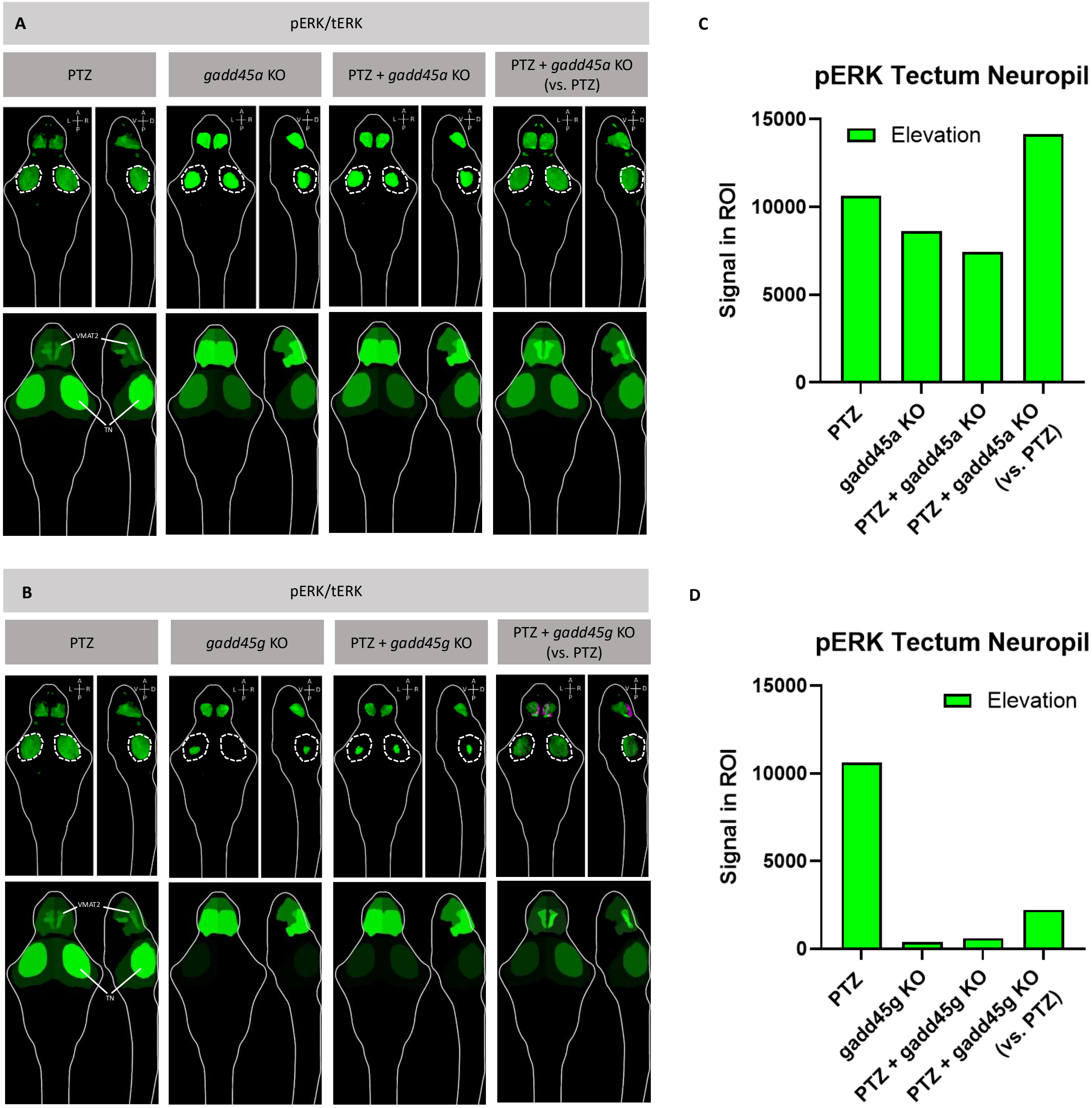


**Supplementary Figure 10.**

(**A-B**) pErk/tErk staining and analysis revealed changes in basal brain activity following various treatment conditions. Green: elevation. Magenta: reduction. PTZ: 1-hr PTZ treatment at 5 dpf; analysis was conducted by comparing to DMSO control; re-adaptation of Figure 3B. *gadd45a/g* KO: F0 CRISPR/Cas9 double-KO of *gadd45aa* and *gadd45ab* (*gadd45a* KO) (C) or *gadd45ga* and *gadd45gb* (*gadd45g* KO) (D); analysis was conducted by comparing to wild-type (WT) control. PTZ + *gadd45q/g* KO: *gadd45a/g* KO larvae exposed to PTZ for 1-hr at 5 dpf; analysis was conducted by comparing to WT control. PTZ + *gadd45a/g* KO (vs. PTZ): *gadd45a/g* KO larvae exposed to PTZ for 1-hr at 5 dpf; analysis was conducted by comparing to WT control exposed to PTZ for 1-hr at 5 dpf. Brain regions are identified based on alignment with the zBrain atlas following brain registration. The top differentially regulated brain regions are labeled in the images. Raw analysis results for all differentially regulated brain regions and their signal intensities in images shown in Supplementary Figure 9 are summarized in Supplementary Data 5. TN: tectum neuropil. VMAT2: vmat2 cluster (telencephalon). (**C-D**) Bar plots showing summed pERK signal changes in the tectum neuropil (white dashed circles in Supplementary Figures 10A & 10B) following

PTZ exposure, *gadd45a/g* KO, combined PTZ + *gadd45a/g* KO, and PTZ + *gadd45a/g* KO relative to PTZ alone. Positive values indicate elevation and negative values indicate reduction in pERK signal. The analysis reveals PTZ-induced increases in brain activity (pERK signal) (PTZ) that are partially reversed by *gadd45a/g* knockout (PTZ + *gadd45a/g* KO).

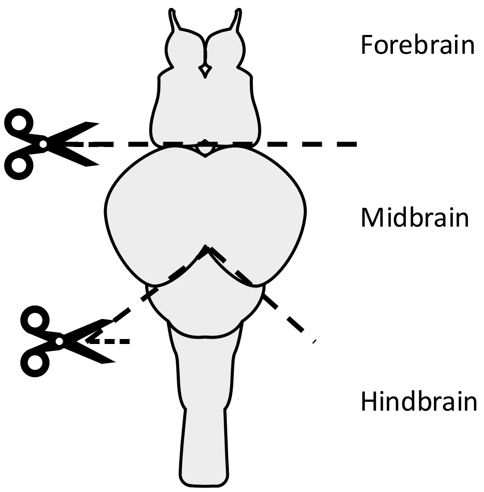


**Supplementary Figure 11.**

Cutting locations for dissecting the larvae forebrain, midbrain, and hindbrain regions.

### SUPPLEMENTARY TABLE LEGENDS

**Supplementary Table 1.**

gRNA sequences.

**Supplementary Table 2.**

Number of larvae imaged in each experimental group.

**Supplementary Table 3.**

qPCR primer sequences.

**Supplementary Table 4.**

Hybridization chain reaction (HCR) probe design.

### SUPPLEMENTARY DATA LEGENDS

**Supplementary Data 1.**

Raw analysis results for images shown in Figure 1, listing all differentially regulated brain regions and their respective signal intensities. Green-colored tables show brain regions with elevated activity while magenta-colored tables show brain regions with reduced brain activity compared to the controls.

**Supplementary Table 2.**

Raw analysis results for images shown in Figure 2, listing all differentially regulated brain regions and their respective signal intensities. Green-colored tables show brain regions with elevated activity while magenta-colored tables show brain regions with reduced brain activity compared to the controls.

**Supplementary Table 3.**

Raw analysis results for images shown in Figure 3, listing all differentially regulated brain regions and their respective signal intensities. Green-colored tables show brain regions with elevated activity while magenta-colored tables show brain regions with reduced brain activity compared to the controls.

**Supplementary Table 4.**

Raw analysis results for images shown in Figure 4, listing all differentially regulated brain regions and their respective signal intensities. Green-colored tables show brain regions with elevated activity while magenta-colored tables show brain regions with reduced brain activity compared to the controls.

**Supplementary Table 5.**

Raw analysis results for images shown in Supplementary Figure 9, listing all differentially regulated brain regions and their respective signal intensities. Green-colored tables show brain regions with elevated activity while magenta-colored tables show brain regions with reduced brain activity compared to the controls.
